# Type I interferons induced upon respiratory viral infection impair lung metastatic initiation

**DOI:** 10.1101/2024.12.23.630113

**Authors:** Ana Farias, Victoria Bridgeman, Felipe S. Rodrigues, Amber Owen, Stefanie Ruhland, Rute Ferreira, Matthias Mack, Ilaria Malanchi, Cecilia Johansson

## Abstract

Invasive breast cancer accounts for 7% of all cancer-related deaths, with the lungs being a common site of metastases. At the same time, lower respiratory tract infections are a common cause of morbidity and mortality worldwide. Acute viral respiratory infections induce transitional changes in the lung; however, the impact of these changes on metastasis initiation and cancer progression remains unclear. Using primary murine MMTV-PyMT breast cancer cells in an experimental lung metastasis model, we show that changes induced by respiratory syncytial virus (RSV) infection impair tumor cell seeding and early establishment in the lung, resulting in lower number of metastatic nodules. Furthermore, we demonstrate that this reduction of metastases is due to alterations in the lung environment mediated by type I interferons (IFNs) that are produced in response to RSV infection. Consistent with that notion, intranasal administration of recombinant IFN-α recapitulates the anti-tumor effect of RSV infection. Type I IFNs change the lung cellular composition and induce an Interferon Stimulated Gene (ISG) driven response, creating an alveolar environment that is less supportive of tumor cell growth. Indeed, epithelial cells from mice infected with RSV or intranasally exposed to IFN-α, are less supportive of tumor cell growth *ex vivo*. Altogether, our results suggest that type I IFNs induced by infection with some respiratory viruses perturb the lungs and consequently interfere with the ability of tumor cells to successfully initiate metastatic colonization.

**Significance:** Women diagnosed with metastatic breast cancer have a low survival rate. The lungs are a common metastatic site and are constantly exposed to viral pathogens, such as coronavirus, RSV and influenza virus. Thus, breast cancer and respiratory virus infection are likely to co-occur, but their interplay remains unclear. We show that type I interferons (IFNs), induced upon viral infection impair metastatic cancer cell seeding of mouse lungs. This is potentially via an effect of IFNs on lung epithelial cells, which become less supportive of early tumor cell proliferation. These findings indicate that viral infections and type I IFNs can alter the lung environment and impair implantation of metastatic cells, which could be explored to improve future cancer treatments.

## Introduction

The lungs are anatomically positioned at the body-environmental interface where they play an essential role in gas exchange. The constant flow of air into the lower airways constantly exposes the lung cells to microbes, allergens, noxious gases and pollution particles. To prevent overexuberant immune response, the lung environment has a high threshold for immune cell activation(1). This provides an optimal niche for distant cancer cells to successfully start metastases, making the lungs the second most frequent site of metastasis for many malignancies, including colorectal cancer, head and neck cancer, breast cancer and melanoma(2). According to the World Health Organization (WHO), cancer is the second leading cause of death globally(3), with invasive breast cancer being the leading cause of cancer deaths among women(4).

Lower respiratory tract infections caused by both viral and bacterial pathogens are associated with high morbidity and mortality rates worldwide(5). Respiratory viruses, such as coronavirus, influenza virus and respiratory syncytial virus (RSV) commonly cause infections in the upper respiratory tract that can progress to the lower tract and result in severe bronchiolitis or pneumonia(6). Seasonal vaccines for influenza virus and coronavirus are available worldwide for high-risk populations, including the immunosuppressed and the elderly, and there are recently-approved vaccines against RSV(7). RSV infection is often associated with childhood bronchiolitis. However, the elderly and immunocompromised are at high risk of developing severe disease and show increased mortality rates(8). Recently, it has been shown that RSV infections in older adults result in increased admissions to intensive care units compared to influenza virus infections(9).

Lung metastases and lower respiratory infections both have an impact on the lung environment and they can co-occur, yet the interplay between them remains unexplored. An association between pneumonia and bronchiolitis and higher risk of lung cancer has been suggested(10,11). However, these epidemiological findings have been based on self-reported pneumonia and primary lung cancer and do not consider the timing, number and types of infections or the stage of lung cancer. Thus, efforts are needed to decipher how changes induced in the lungs by viral infections and lung malignancies impact on the outcome of disease.

Many respiratory viruses infect lung epithelial cells and quickly induce a pro-inflammatory response that results in the activation of resident cells and the recruitment of innate immune cells(12,13). This rapid innate immune response limits viral replication and orchestrates the adaptive immune response, generating cytotoxic CD8^+^ T cells and a humoral response, important for viral clearance and for protection against subsequent infections(14). These immune responses change the lung environment and could therefore have an impact on lung cancer initiation, progression, as well as on metastatic spread to the lungs from different primary tumors. Cuff et al., showed that influenza virus infection enhances the development of experimental lung metastasis in the B16 melanoma model by activation of bystander CD8^+^ T cells not specific for the tumor antigens(15). However, Newman et al., described that a productive influenza virus infection leads to a potent immune response that reduces the number of B16 metastatic nodules in the lungs(16). Although both studies used the same metastatic cancer model, infections were conducted at different times prior or after tumor cell injection, suggesting that the interplay between these diseases is temporally dynamic. Deciphering the mechanisms underlying the different outcomes in these studies could provide insights on how to improve cancer treatments.

Type I interferons (IFNs) comprise IFN-β and many IFN-α subtypes and are important anti- viral cytokines(13). During RSV infection, alveolar macrophages (AMs) are the main producers of type I IFNs, responding to the virus via mitochondrial antiviral signaling protein (MAVS)-coupled retinoic acid-inducible gene I (RIG-I)-like receptors (RLRs)(17). Type I IFNs inhibit viral replication but are also involved in the recruitment of antiviral monocytes and activation of immune cells, which all together control infection and disease severity(12,17). Due to the ability of type I IFNs to modulate innate immune responses and to efficiently orchestrate the adaptive immune responses, their role in cancer immunosurveillance and their anti-tumoral and pro-tumoral effects have been extensively studied(18–22). However, the role of type I IFNs induced by respiratory viral infections in metastasis initiation and progression remains unclear.

We hypothesized that the early type I IFN-driven proinflammatory response induced by RSV infection influences the lung seeding and growth of breast cancer metastases. We therefore studied the ability of intravenously inoculated mouse mammary tumor virus-polyoma middle tumor-antigen cells (MMTV-PyMT cells) to seed and grow in the lungs of RSV-infected mice. We show that RSV infection a day before cancer cell injection reduces the seeding of the MMTV-PyMT cells in the lung, resulting in lower number of metastatic nodules. Moreover, we demonstrate that this reduction is due to type I IFNs changing the lung environment, generating less supportive conditions for metastatic initiation.

## Results

### RSV infection reduces the number of metastatic nodules in the lungs

To investigate if RSV infection impacts the ability of metastatic cells to seed the lungs, survive and proliferate, we synchronized cancer cell seeding by using an experimental lung metastasis model. Primary breast carcinoma cells (MMTV-PyMT cells) were injected intravenously (i.v.) into FVB/N or C57BL/6J mice 24h after intranasal (i.n.) infection with RSV (Fig. 1A). Tumor burden was assessed 28 days after cell administration by histological analysis and macroscopic nodule quantification. Interestingly, we found that FVB/N and C57BL/6J mice displayed lower number of metastatic nodules in the lungs when they were previously infected with RSV (Fig. 1B-D and Supp. Fig. 1). However, the metastatic nodules that did develop in RSV infected showed a similar size distribution to those in control mock-infected (PBS instillation) mice (Fig. 1B-D). Similar results were found in BALB/c mice using 4T1 breast cancer cells (Supp. Fig. 2). These data suggest that RSV infection inhibits metastatic lung colonization but that, once established, tumor cells can grow unimpeded.

**Figure 1.**
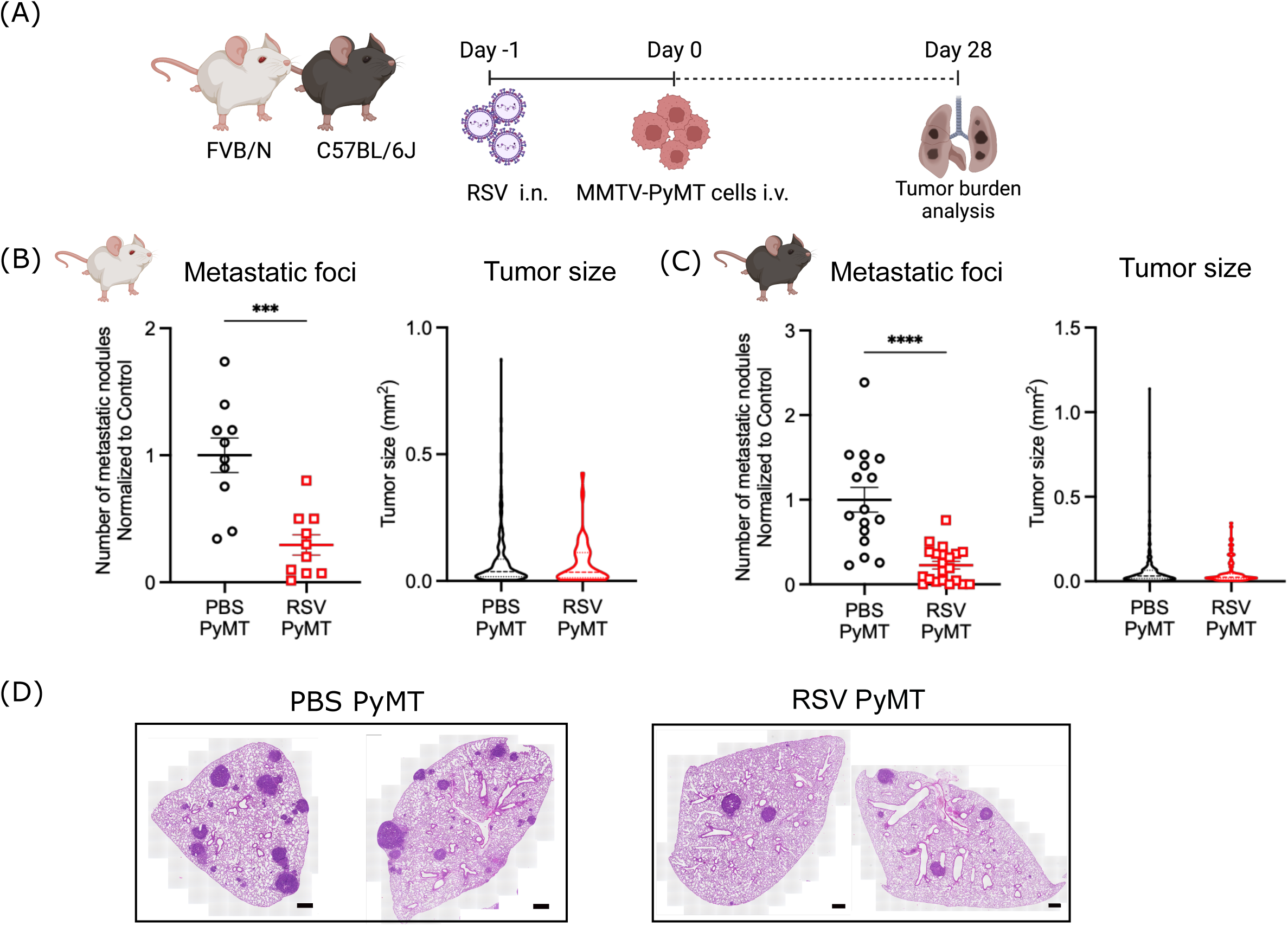
RSV infection impairs the development of metastases in the lungs. (A) Experimental setup. (B) FVB/N or (C) C57BL/6J mice were intranasally (i.n.) infected with RSV (RSV PyMT) or treated with PBS (PBS PyMT). A day later, 3x10^5^ MMTV-PyMT cells were intravenously (i.v.) inoculated. Tumor burden was analyzed 28 days post tumor cells inoculation. To assess tumor burden, 3 levels at least 150 mm apart of all lobes were analyzed by H&E staining. Total number of metastatic nodules in (B) and (C) were normalized to the average of the uninfected (PBS) group in each independent experiment. Tumor size shows the size of all tumors detected in the group. (D) Representative H&E-stained sections of the lungs of FVB/N mice at 28 days post cell injection, scale bar 500 µm. Data for FVB/N are pooled from two independent experiments presented as the mean +/- SEM of 10 mice per group. Data for C57BL/6J are pooled from three independent experiments presented as the mean +/- SEM of 16 mice for mock infected and 23 mice for the RSV infected group. Student’s t-test was performed. Only statistically significant differences are shown; *** p<0.001; **** p<0.0001.

### The presence of metastatic cells in the lung does not influence the overall response to RSV infection

Using FVB/N mice, we also assessed if the presence of lung metastases influences the course of RSV infection (Fig 2A). FVB/N mice developed severe disease during RSV infection (Fig 2B), with increased weight loss compared to that which we have previously reported in C57BL/6J mice(23). Interestingly, administration of MMTV-PyMT cells a day after infection did not alter subsequent weight loss (Fig. 2B) or viral load (Fig. 2C). The total number of cells, mostly CD45^+^, recovered from the lungs and the bronchoalveolar lavage (BAL) was quantified at different times post infection (p.i.) (Fig. 2D-F). RSV infection led to increased abundance of cells that could be recovered from the lungs and BAL at days 4 and 8 p.i., respectively. Interestingly, increased number of cells were detected at 18h and 3 days post cell administration (d2 and d4 p.i.) in all mice bearing MMTV-PyMT cells irrespective of RSV infection (Fig. 2E).

**Figure 2.**
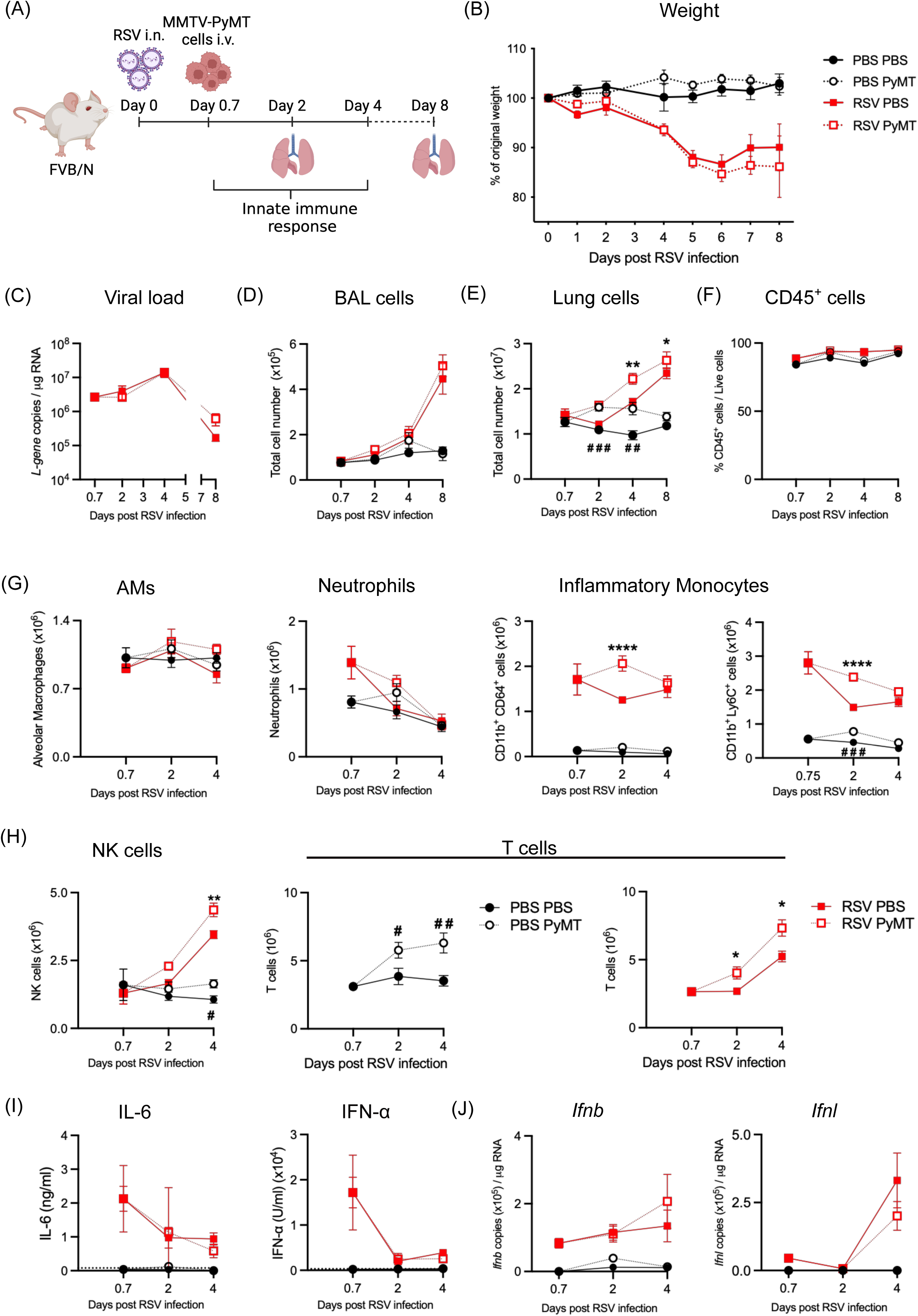
Inoculation of tumor cells a day after RSV infection does not alter the anti-viral immune response. (A) FVB/N mice were intranasally (i.n.) infected with RSV or treated with PBS. A day later, 3x10^5^ MMTV-PyMT cells were intravenously (i.v.) administered (PBS PyMT and RSV PyMT). (B) Disease severity was assessed by weight loss plotted as % of original weight. (C) Viral load was quantified by RT-qPCR at different time points post infection from RNA isolated from lung tissue. Cells in the airways (BAL; D) and lungs (E) were enumerated. (F) Percentage of CD45^+^ cells recovered from the lungs quantified by flow cytometry. (G) Numbers of alveolar macrophages (AMs), neutrophils, inflammatory monocytes (gated as CD11b^hi^ CD64^hi^ or Ly6C^+^CD11b^hi^), (H) NK cells and T cells present in the lungs at different times post infection were analyzed by flow cytometry. (I) Levels of IL-6 and IFN-a in BAL were quantified by ELISA. (J) Copy numbers of mRNA *Ifnb* and *Ifnl* were quantified by RT-qPCR. Weight loss data (B) are pooled from 3 independent experiments and shown as mean +/- SEM of n=13 for both infected groups and n=12 for the MMTV-PYMT mock infected group. Data for the PBS group are pooled from two experiments and shows as mean +/- SEM of n=8. A two-way ANOVA, mixed-effect analysis was performed to compare weight loss after infection followed by Tukey’s post hoc test. (C-J) Data are pooled from two independent experiments presented as the mean +/- SEM; for day 0.7; n= 6 mice in the PBS group and n= 7 in the RSV group, for day 2; n= 9 mice per group. Days 4 and 8; n= 8 mice per group. One-way ANOVA was performed to compare the PBS and PyMT mice (#) and the RSV and RSV PyMT mice (*) at each time point. Only statistically significant differences are shown; * p<0.05, ** p<0.01, ****p<0.0001; # p<0.05, # # p<0.01, # # # p<0.001.

To further characterize the immune response induced by RSV in the presence of tumor cells, alveolar macrophages, neutrophils, inflammatory monocytes, NK cells and T cells were quantified at different time points by flow cytometry (gating strategy Supp. Fig. 3) in the lungs (Fig. 2G-H) and in the BAL (Supp. Fig. 4A). RSV infection in FVB/N mice did not alter the number of alveolar macrophages (AMs; Fig. 2G and Supp. Fig. 4A). However, the infection resulted in recruitment of neutrophils, with higher numbers of these cells detected 18h post infection (Fig. 2G,(24)). Similarly, inflammatory monocytes (gated as Ly6G^-^ SiglecF^-^ CD11b^+^CD64^+^ cells or Ly6G^-^ SiglecF^-^ CD11b^+^ Ly6C^+^ cells) were detected as early as 18h p.i., with higher numbers in the PyMT-bearing mice at day 2 post-RSV infection (Fig. 2G). We also detected an increase in the abundance of NK cells, peaking at 4 days p.i., with higher numbers in the RSV-infected mice that also received PyMT cells (Fig. 2H). A transient increase in T cells was observed in the lungs of PyMT-bearing mice at day 2 and 4 p.i. but only at day 4 p.i. in RSV-infected animals that did not receive tumor cells (Fig. 2H).

We next studied the expression of immune mediators triggered by RSV infection in the presence of tumor cells. In infected groups, similar levels of IL-6 and IFN-α were detected in the BAL fluid by ELISA (Fig. 2I) and in lung tissue by analysis of gene transcripts (Supp. Fig. 4B). Furthermore, levels of mRNA encoding Ifnb or Ifnl (Fig. 2J) and Il1b, Ccl2 or Cxcl1 (Supp Fig. 4B) in lung tissue were similar across all the RSV-infected groups irrespective of tumor cell injection. We also studied the expression of interferon stimulated genes (ISGs), including Pkr, Viperin, Cxcl10, Mx1 and Oas1 (Supp Fig. 4C). The same ISG expression profile was induced after RSV infection regardless of the presence of tumor cells (Supp. Fig. 4B-C). These data suggest that the overall innate immune response against RSV infection is largely unaffected by the presence of tumor cells in the lungs, with only a transient increase of inflammatory monocytes, NK cells and T cells in mice in which infection proceeds concomitantly with the presence of tumor cells.

### Neutrophils, monocytes or NK cells are not essential to impair tumor cell metastases during RSV infection

Type I IFNs are induced by alveolar macrophages upon detection of RSV(17). Interestingly, type I IFNs can induce an anti-tumor response by activating innate immune cells, including neutrophils, monocytes and NK cells(18,22,25–28). To assess a potential role of neutrophils in the anti-tumoral response we used anti-Ly6G mediated neutrophil depletion during RSV infection (Fig. 3A and Supp. Fig. 5A). Interestingly, no differences in numbers of metastatic foci or tumor size were detected between infected mice with or without neutrophils (Fig. 3B). Neutrophil depletion also had no impact on RSV-induced disease severity (Supp. Fig. 5B). We have previously shown that mice lacking TRIF and MyD88 adaptor proteins are unable to recruit neutrophils to the lungs during RSV infection(24). MyD88/TRIF deficient mice were infected with RSV and inoculated with tumor cells i.v. a day later (Supp. Fig. 5C). Analysis of tumor burden showed that RSV-infected Myd88/Trif^-/-^ mice and wildtype mice displayed a similar decrease of metastatic nodules in their lungs after RSV infection (Supp. Fig. 5D). Together, these data suggest that RSV-induced recruitment of neutrophils to the lungs does not have an essential role in reducing lung metastatic colonization.

**Figure 3.**
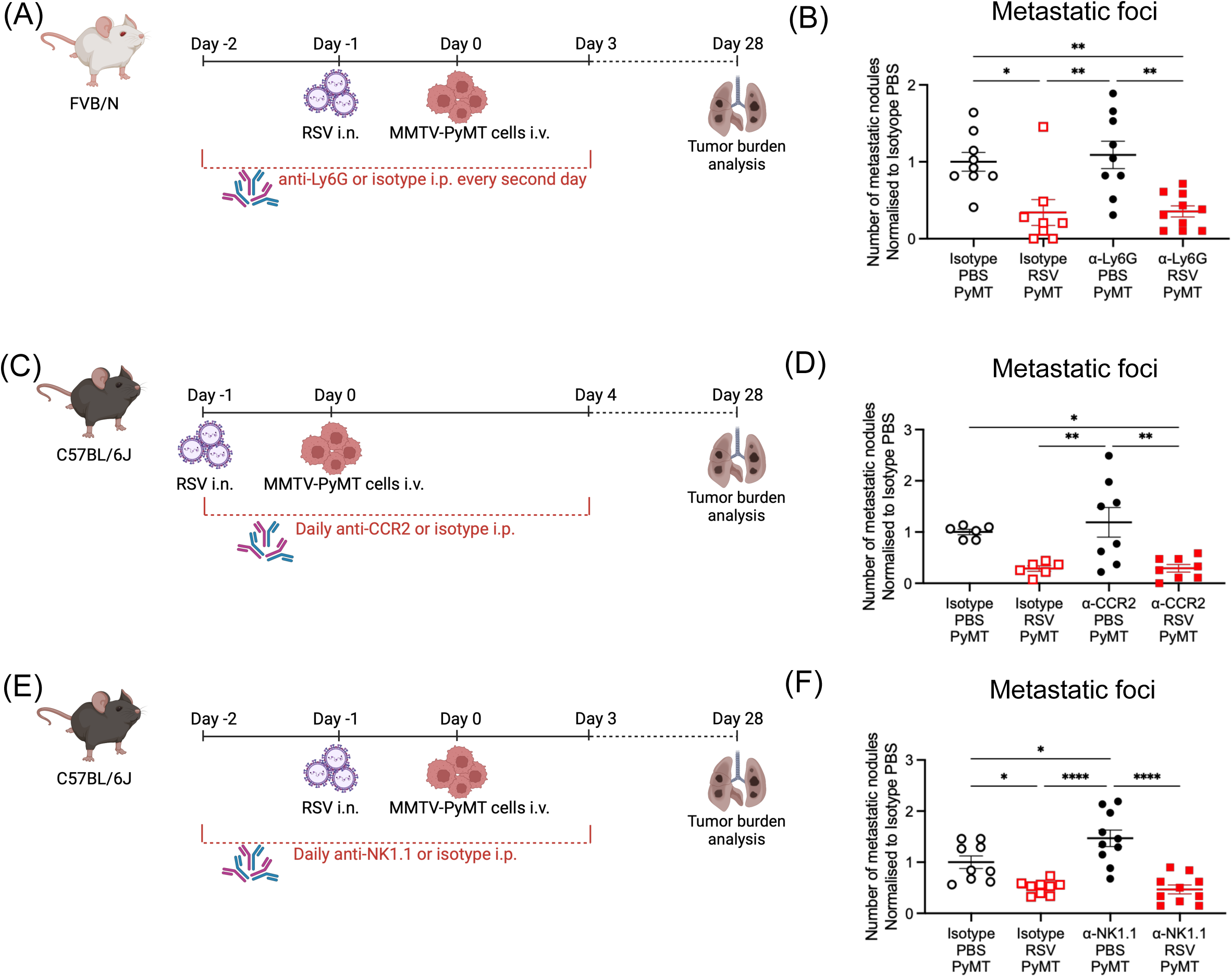
Depletion of a single immune cell population during RSV infection does not identify a specific cell type that impairs metastatic burden. (A) FVB/N mice were treated with 150µg of anti-Ly6G or isotype intraperitoneally (i.p.) every second day starting a day prior to RSV infection up to day 4 p.i.. A day after RSV infection, MMTV-PyMT cells were administered i.v.. (B) After 28 days, tumour burden was assessed by H&E staining. Data are pooled from two independent experiments presented as the mean +/- SEM of 9 mice per group in the PBS group and 8 or 10 mice in RSV infected mice treated with isotype or anti- Ly6G, respectively. (C) C57BL/6J mice were treated daily with 20 µg anti-CCR2 or isotype from the day of RSV infection until 5 days p.i.. MMTV-PyMT cells were injected i.v. a day after RSV infection as previously described. (D) Metastatic burden was evaluated 28 days after tumor cell injection. Data are pooled from two independent experiments presented as the mean +/- SEM of 6 mice per group in the isotype treated groups and 8 mice in the anti- CCR2 groups. (E) To deplete lung NK cells, 100 µg of anti-NK1.1 was administered i.n. a day before infection. Depletion was maintained from the day prior to infection until day 4 p.i. by daily injection of 150 µg of anti-NK1.1. (F) Lung metastatic burden was analyzed 28 days later. Data are pooled from two independent experiments presented as the mean +/- SEM of 9 mice per group in the isotype treated groups and 10 mice in the anti-NK1.1 groups. The relevant isotype was used in the control groups. One-way ANOVA was performed to compare all groups followed by Tukey’s post hoc test. Only statistically significant differences are shown; * p<0.05, ** p<0.01, ****p<0.0001.

We next studied the role of monocytes in the reduced metastatic burden after RSV infection. Monocytopenia was achieved using anti-CCR2 treatment(29) from the day of infection until day 5 p.i. (Fig. 3C and Supp. Fig. 5E). Disease severity caused by RSV infection was not altered in the absence of monocytes (Supp. Fig. 5F). Notably, lack of monocytes did not impact the number of metastatic foci as this was equally reduced in isotype-matched control vs. anti-CCR2 treated mice after RSV infection (Fig. 3D).

RSV infection also cause an increase in the number of NK cells in the lungs, which is further potentiated by the presence of cancer cells (Fig. 2H). We therefore depleted NK cells using anti-NK1.1 antibody before and during RSV infection. This did not impact weight loss caused by the infection (Fig. 3E and Supp. Fig. 5G and H) and, strikingly, did not curtail the inhibitory effect of the infection on metastatic establishment (Fig. 3F). However, as expected, NK cell depletion in mock-infected mice led to an increase in lung metastases (Fig. 3F). These results indicate that NK cells are not necessary for the reduction in lung metastatic initiation caused by RSV infection. Altogether, our results show that despite the RSV-dependent increase in neutrophils, monocytes and NK cells, each of those immune cells do not explain the virus-induced anti-tumoral response.

### The T cell response during RSV infection does not play an essential role in decreasing the metastatic burden

We next studied T cell responses at day 8 post RSV infection (Fig. 4A). At this timepoint, we noticed clear differences in T cell number between infected and uninfected mice, yet these numbers were unaltered by PyMT inoculation (Fig 4B-D, Supp. Fig. 6A). Indeed, both RSV-infected tumor-free and tumor-inoculated groups displayed the same number of total T cells, specifically CD8^+^ T cells, in the lungs (Fig. 4B-C), as well as CD4^+^ T cells and CD8^+^ T cells in the BAL (Supp. Fig. 6A). Activation of T cells, as determined by expression of CD69 and PD1, was also similar in all infected mice irrespective of inoculation with tumor cells (Fig. 4C and Supp. Fig. 6A). Further, levels of IFN-γ and Granzyme B detected in BAL fluid were similar across infected groups (Fig. 4D). Overall, these results suggest that the T cell response induced by RSV infection is not significantly affected by the presence of MMTV-PyMT cells in the lungs.

**Figure 4.**
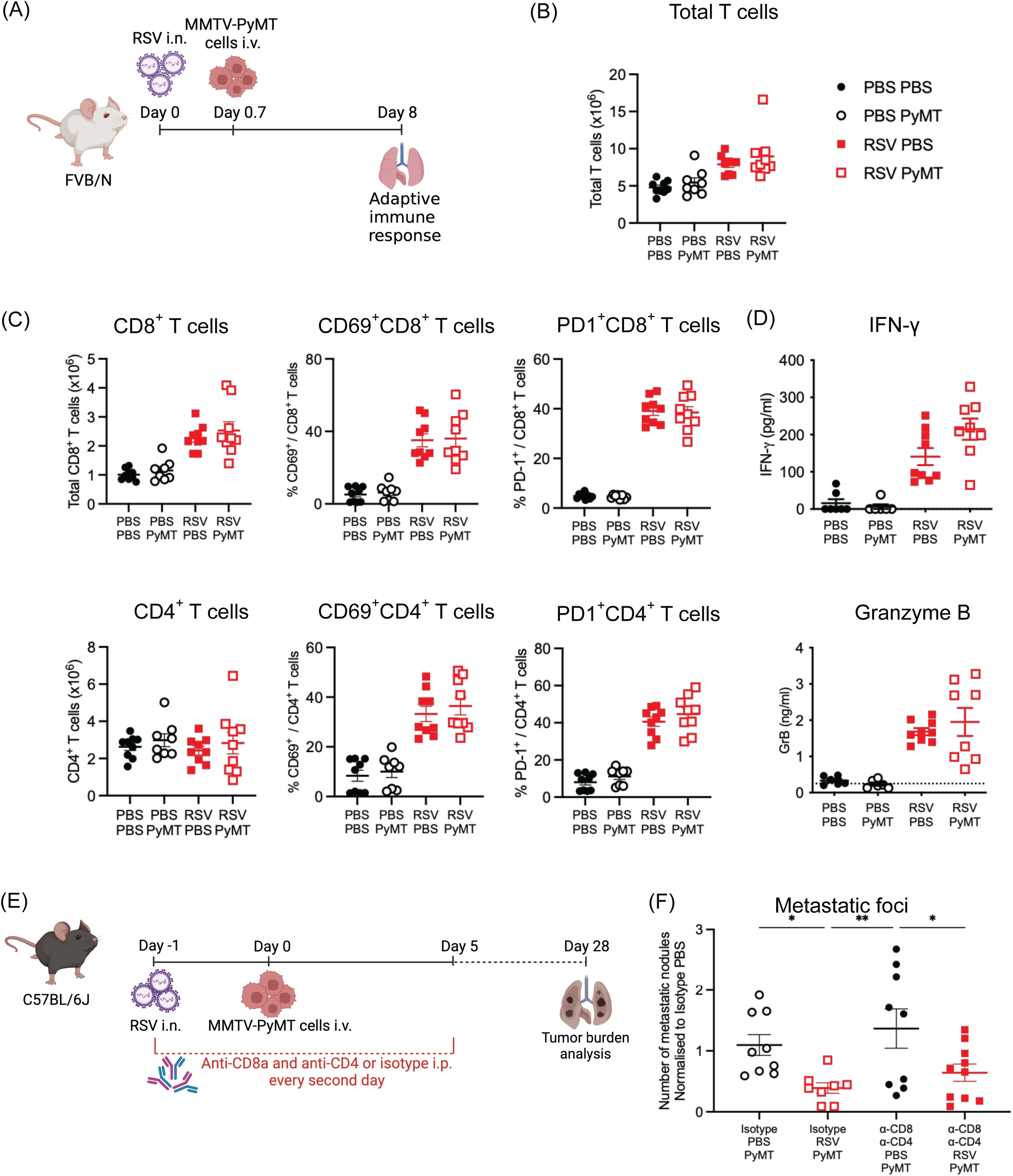
The T cell response elicited by RSV infection is not altered in MMTV-PyMT bearing mice. (A) FVB/N mice were infected i.n. with RSV or mock-infected with PBS. A day later, 3x10^5^ MMTV-PyMT cells were inoculated i.v.. (B) Number of T cells (CD3^+^), (C) CD8^+^ T cells and CD4^+^ T cells as well as activated T cells measured as CD69^+^ or PD1^+^ T cells were quantified in lungs 8 days p.i., following the gating strategy shown in Supp. Fig. 3B. (D) At 8 days p.i.. IFN-γ and Granzyme-B in BAL were quantified by ELISA. Data are pooled from two independent experiments with 8 mice per group. (E) CD4^+^ and CD8^+^ T cells were depleted during RSV infection by i.p. administration of anti-CD4 and anti-CD8 or isotype control every second day starting the day of infection until day 6 p.i.. MMTV-PyMT cells were injected i.v. a day after RSV infection. One-way ANOVA was performed to compare infected groups followed by Tukey’s post hoc test. No significant differences were detected. (F) Tumor burden was assessed by H&E quantification 28 days after tumor cell administration. Data are pooled from two independent experiments presented as the mean +/- SEM of 9 mice per group in the PBS group and 8 or 10 mice in RSV infected mice treated with isotype or anti-CD4 and anti-CD8, respectively. One-way ANOVA was performed to compare all groups followed by Tukey’s post hoc test. Only statistically significant differences are shown; * p<0.05, ** p<0.01: ***p<0.001

To formally evaluate the role of T cells on the anti-tumoral effect induced by RSV infection, CD8^+^ and CD4^+^ T cells were depleted with antibodies (Fig. 4E and Supp. Fig. 6B). As previously shown(30), depletion of T cells decreases the weight loss seen at day 5-7 p.i. (Supp. Fig. 6C), which is due to immunopathology. However, regardless of the presence of T cells, RSV infection still led to a lower number of metastatic nodules compared to uninfected mice (Fig. 4F). Thus, T cells are also dispensable for the reduced metastatic burden induced by RSV infection.

### Type I interferons recapitulate the anti-tumoral effect observed after RSV infection

Type I interferons (IFN) are key anti-viral cytokines released early upon RSV infection and responsible for inhibiting viral replication and orchestrating the innate immune response by signaling through the IFNα/β receptor (IFNAR)(12). We first analyzed how IFN-α changes the lung environment within 18h of airway exposure (Supp. Fig. 7A). Administration of IFN-α i.n. to C57BL/6J mice or to FVB/N mice resulted in a transient inflammatory response characterized by activation of resident neutrophils (determined as CD64^+^ neutrophils) and recruitment of inflammatory monocytes (Supp. Fig. 7B-D,(12)). Expression of several chemokines such as Ccl2 and Cxcl1 and cytokines including ll1b and Il6 and ISGs was also increased at this time point (Supp. Fig. 7E-F). Thus, type I IFNs are sufficient to remodel the lung environment 18h after inhalation, the timepoint when tumor cells are injected i.v. in our model. To note, the presence of MMTV-PyMT cells, does not impact the overall lung immune response induced by IFN-α (Supp. Fig. 8).

To determine if type I IFNs phenocopy the anti-tumoral effect of RSV infection, we administered two doses of recombinant IFN-α (IFN-α) intranasally to C57BL/6J mice, 24 and 18h prior to cancer cells injection and assessed tumor burden 28 days later (Fig. 5A). Remarkably, compared to the control group, IFN-α exposure resulted in reduced number of metastatic nodules in the lung, to a similar extent as RSV infection (Fig. 5B). Furthermore, UV-inactivated RSV, which does not induce a productive infection and therefore does not induce type I IFNs(31), did not offer protection from metastatic burden (Fig. 5B). Thus, early administration of type I IFNs, like RSV infection, has an anti-metastatic effect.

**Figure 5.**
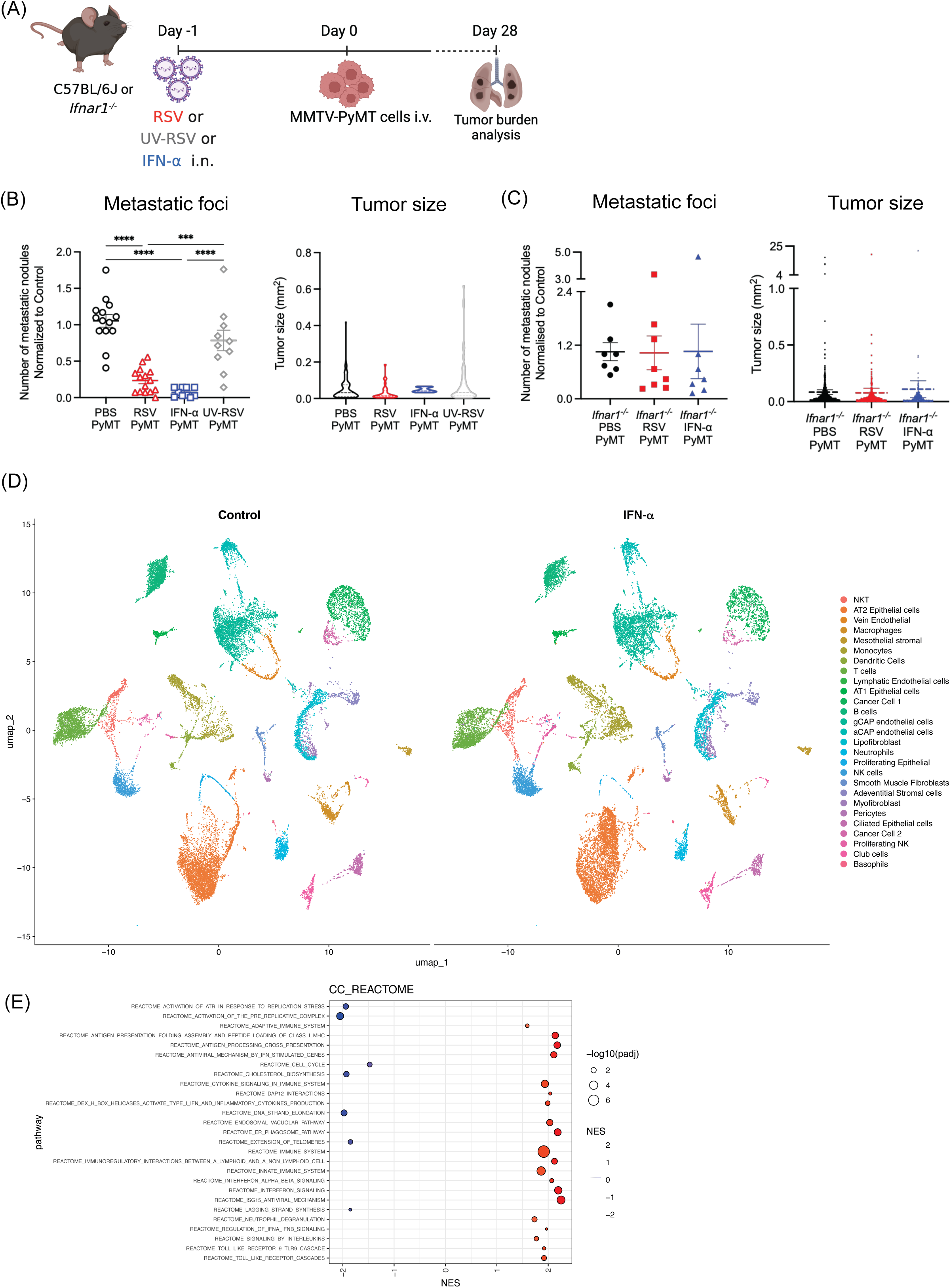
Lower number of metastatic nodules in the lung following RSV infection is dependent on type I IFN receptor signaling. (A) Experimental setup. C57BL/6J mice were intranasally mock infected (PBS PyMT) or infected with RSV (RSV PyMT) or exposed to two doses of 500 ng IFN-α (18h apart, with the second dose at least 4h before tumor cell injection; IFN-α PyMT) or exposed to UV inactivated RSV (UV-RSV PyMT). MMTV-PyMT cells were injected i.v. a day later. (B) Number and size of metastatic nodules was determined by H&E staining 28 days after tumor cell injection. (C) *Ifnar1^-/-^* mice were either mock infected (PBS), infected with RSV or exposed to two doses of 500 ng IFN-α followed by i.v. inoculation of MMTV-PyMT cells. Number and size of metastatic nodules was analyzed 28 days later by H&E staining of 3 levels of each lobe, total number of metastatic tumors was normalized to the average of the uninfected group in each independent experiment. Data from tumor size quantification were pooled from all detectable tumors in all mice in each group. Data are presented as the mean +/- SEM pooled from two independent experiments, for C57BL/6J mice n= 14 for PBS PyMT, n=15 for RSV PyMT, n=10 for IFN-α PyMT and n=10 for UV-RSV PyMT. For *Ifnar1^-/-^* mice n=7 for PBS PyMT and IFN-α PyMT and n=8 for RSV PyMT. One-way ANOVA test followed by Tukey’s post hoc test was performed to compare all groups. Only statistically significant differences are shown; ***p<0.001; ****p<0.0001. (D) Experimental setup. C57BL/6J mice were intranasally exposed to PBS (Control) or exposed to 1mg IFN-α (IFN-α) , MMTV-PyMT cells were injected i.v. 18 hours later. Lungs were collected 24 hours after i.v. (E) UMAP plot of scRNAseq data from lungs of mice, Control (PBS treated) and IFN-α treated (F) fGsea analysis of Cancer Cell Reactome pathways only showing the top statistically significant pathways with a Normalized Enrichment Score (NES) >1 or < -1.

To assess if RSV-induced type I IFNs acting on host cells are necessary for metastatic reduction, we also infected mice lacking the type I IFN receptor (Ifnar1^-/-^mice; Fig. 5A). Notably, RSV infection in Ifnar1^-/-^ mice did not decrease numbers of metastatic nodules or size compared to the uninfected group (Fig. 5C). To further assess if IFN-α impacts tumor cells directly, Ifnar1^-/-^mice were treated with IFN-α to create a setting in which only the tumor cells can respond to the cytokine. IFN-α administration to Ifnar1^-/-^ mice did not change the number of metastatic nodules or size compared to the mock treated group (Fig. 5C). Overall, these data suggest that IFN-α induces changes in the lung environment rather than acting directly on the tumor cells to reduce metastatic initiation.

To further explore how the lung response to IFN-α influences the interaction with the metastatic cells, we performed scRNAseq analysis on lung cells from mice that had been exposed to PBS or IFN-α 18hrs prior to tumor cell injection. The lungs were collected 24hrs after the tumor cells were injected. In order to capture a good representation of the different cell types of the lung we sorted cancer, epithelial, immune and mesenchymal cells using FACS and analyzed them by single cell RNA analysis at a ratio of 1:3:3:3, respectively (Fig. 5D). As predicted, GSEA analysis of the cancer cell cluster showed that cancer cells that were present in the IFN-α pre-treated lung showed a reduction in pathways linked to cell cycle and DNA replication (Fig. 5E). These data further confirmed that none of the immune cells altered their interactions with the cancer cells after IFN-α exposure (Supp. Fig. 9A-C).

### PyMT cell seeding, and early tumor cell growth is blunted after RSV infection or IFN-*α* administration

To study how changes induced by type I IFN exposure of the lungs affect metastatic cells establishment in the tissue we used CellChat to detect changes in interactions coming from lung cells. We first assessed interactions coming from the lung vascular environment to the cancer cells as they enter the tissue via the circulation. Whilst no changes were detected in the interaction between endothelial cells and pericytes, we could observe an increase in their interactions with cancer cells upon IFN-α exposure (Fig. 6A-B). To test whether this could influence the ability of cancer cells to seed the lung, we followed *in vivo* tumor cell extravasation by injecting luciferase-expressing MMTV-PyMT cells into mice after exposure to RSV or IFN-α (Fig. 6C). Bioluminescence was monitored at different time points after cell injection. Notably, RSV infection, as well as IFN-α administration, resulted in a reduced luciferase signal in the lung as early as 3.5h post injection, indicating a reduced ability of circulating cancer cells to seed the lung (Fig. 6D-E). This was confirmed by flow cytometry, using GFP-expressing MMTV-PyMT cells (Supp. Fig. 10). The reduced luciferase signal detected at 2- and 3-days post injection in the control group was very similar in all three experimental groups, suggesting that extravasation of metastatic cells per se is not affected by exposure to RSV or IFN-α (Fig. 6D).

**Figure 6.**
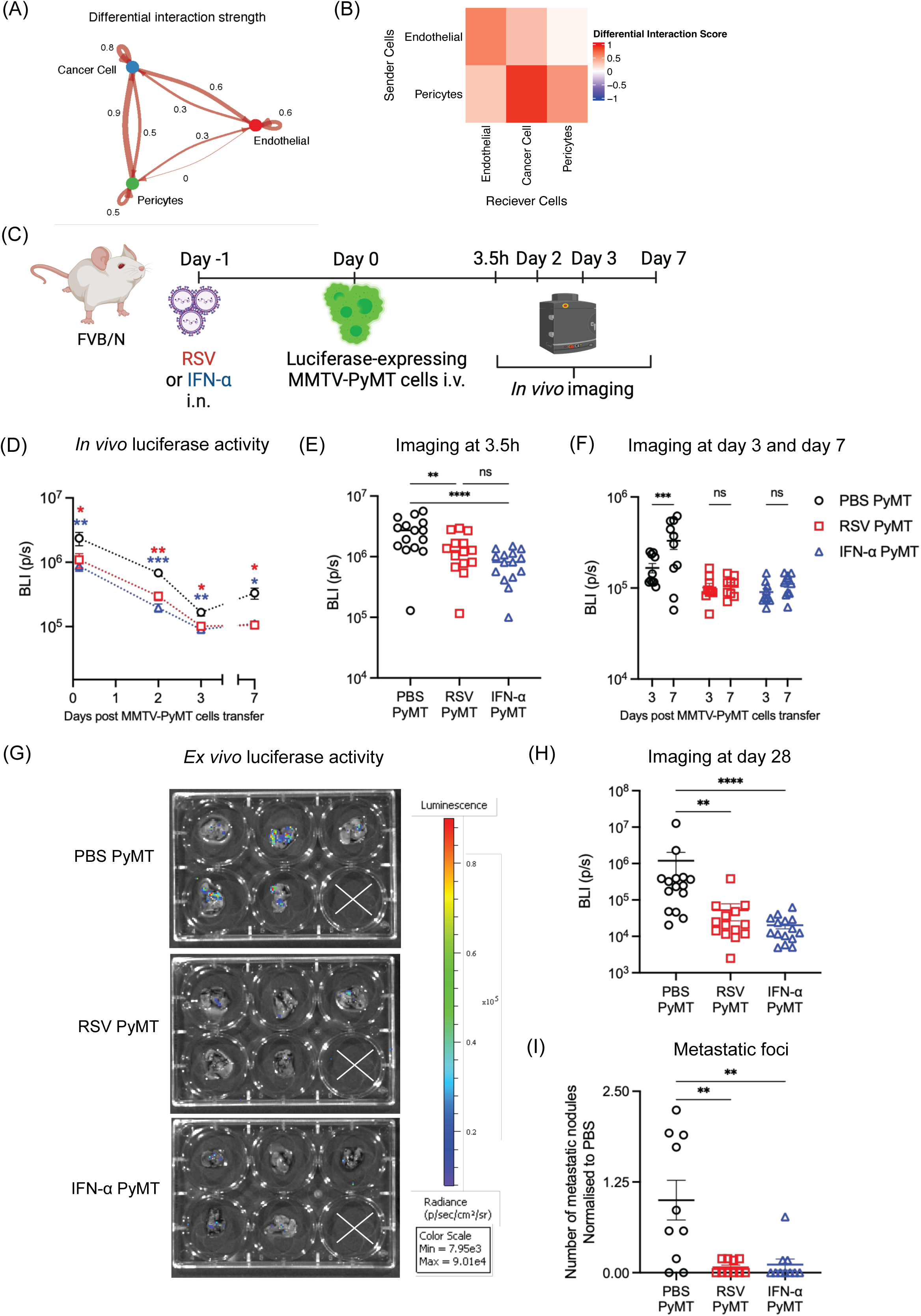
RSV infection inhibits tumor cell seeding in the lung and early cancer cell growth in a type I IFN dependent manner. (A) C57BL/6J mice were intranasally treated with PBS (Control) or one dose of 1ug of IFN-α (IFN-α) and 18h later MMTV-PyMT cells were inoculated i.v. and 24h later the lungs cells were used for scRNA analysis. CellChat analysis was used to create Circle plot of Differential Interaction Score of Cancer Cells with vascular cell components as based on Secreted ligands and Receptor interactions. (B) Heatmap of Differential Interaction Score showing endothelial and pericytes as sender cells and endothelial, cancer cells and pericytes as receivers. (C) FVB/N mice were intranasally treated with PBS (PBS PyMT), one dose of 1ug of IFN-α (IFN-α PyMT) or infected with RSV (RSV PyMT). A day later luciferase expressing MMTV-PyMT cells were inoculated i.v.. (D) At different time points after cell injection, luminescence signal (BLI, photons/second) was quantified using Living Image Software by selecting a ROI of a set size positioned over the thorax of each animal. (E) *in vivo* quantification of luciferase activity 3.5h after MMTV-PyMT cell injection. (F) Comparison of *in vivo* luciferase activity at day 3 and 7 for each experimental group. (G-H) *ex vivo* imaging was performed at day 28 to determine tumor burden, representative images from the different group are shown as well as the quantification of the luciferase activity pooled from three independent experiments, n=15 per group. (I) Number of metastatic nodules was analyzed 28 days later by H&E staining of 3 levels of each lobe, total number of metastatic tumors was normalized to the average of the uninfected group in each independent experiment *in vivo* data are pooled from three independent experiments for day 0 and day 2 (n=15) and two independent experiments for day 3 and 7 (n=10). All data are shown as mean +/- SEM. Statistical significance for the *in vivo* assay was determined by performing a two-way ANOVA with Tukey’s multiple comparisons. For the *ex vivo* readout a Kruskal-Wallis test with Dunn’s multiple comparisons statistical analysis was performed against the PBS group. Data are pooled from two independent experiments, with n=10 mice per group. One-way ANOVA test followed by Tukey’s post hoc test was performed to compare all groups. Only statistically significant differences are shown; *p<0.05; **p<0.01; ***p<0.005; ****p<0.001

The Galectin pathway was major type I IFN induced gene driving the main increased pathway across the different lung cell types, and its activating ligand, Galectin 9 (Lgals9), was strongly increased also in endothelial cells and pericytes (Supp. Fig. 11). Interestingly, Galectin 9 was reported to decrease the ability of cancer cells to extravasate (32). This reduction in the ability of cancer cells to seed on the lung vasculature, alone could explain the reduction in metastatic foci observed upon RSV infected or IFN-α exposure.

Interestingly, in control lungs, we observed an increase in the luciferase signal from day 3 to day 7, indicative of the ability of PyMT cells to initiate early growth post extravasation (Fig. 6F). This increase was not seen in RSV infected or IFN-α exposed mice, pointing at a reduced ability of cancer cells to initiate grow in those conditions (Fig. 6D and F). This resulted in an overall *ex vivo* reduction in bioluminescence at the day 28 end point, which was confirmed to reflect lower tumor burden by histological quantification (Fig. 6G-I).

### Lung epithelial cells from RSV infected or IFN-*α* exposed mice are less supportive of MMTV-PyMT cell growth

Fibroblasts are one of the most dominant tissue cells contributing to the tumor niche, and their early activation is a key factor for metastatic niche initiation of breast cancer cells(33). We and others have shown that also epithelial cells are important cellular components of the metastatic niche in breast cancer lung metastasis(34–37). For example, lung epithelial cells, which rapidly respond to RSV infection(14,38), were shown to directly support cancer cells growth in *ex vivo* 3D co-culture assays(36). Changes in the interaction between mesenchymal and lung epithelial cells were detected upon IFN-α exposure (Fig. 7A-B, Supp. Fig 9D-G), thus, we evaluated if exposure to RSV or IFN-α can affect the interactions between fibroblasts or ATII epithelial cells and the tumor cells that can determine early cancer cell growth. Mice were given either RSV, IFN-α or PBS i.n. and 18h later lung fibroblasts (CD45^-^CD31^-^Epcam^-^) and lung epithelial cells (CD45^-^CD31^-^Epcam^+^) were isolated by FACS (gating strategy Supp. Fig. 12). An Alvetex^TM^ scaffolds co-culture system that mimics the 3D tissue environment was used to culture GFP-expressing MMTV-PyMT cells in presence of the isolated fibroblasts or epithelial cells (Fig. 7C). Tumor cell growth was subsequently determined by the intensity of GFP signal at days 2, 5 and 8. Culture of tumor cells with fibroblasts from PBS-exposed mice induced cell proliferation, detectable as early as day 5. This was not altered in the presence of fibroblasts from mice infected with RSV or treated with IFN-α at any time point (Fig. 7D). Co-cultures with epithelial cells from mock-treated mice showed significantly enhanced PyMT proliferation at day 8. Interestingly, when epithelial cells were isolated from RSV infected or IFN-α exposed lungs, they failed to support cancer cell proliferation to the levels seen in the control group (Fig. 7E). These results demonstrate that epithelial cells from lungs of RSV-infected or type I IFNs exposed mice are less able to support tumor cell proliferation.

**Figure 7.**
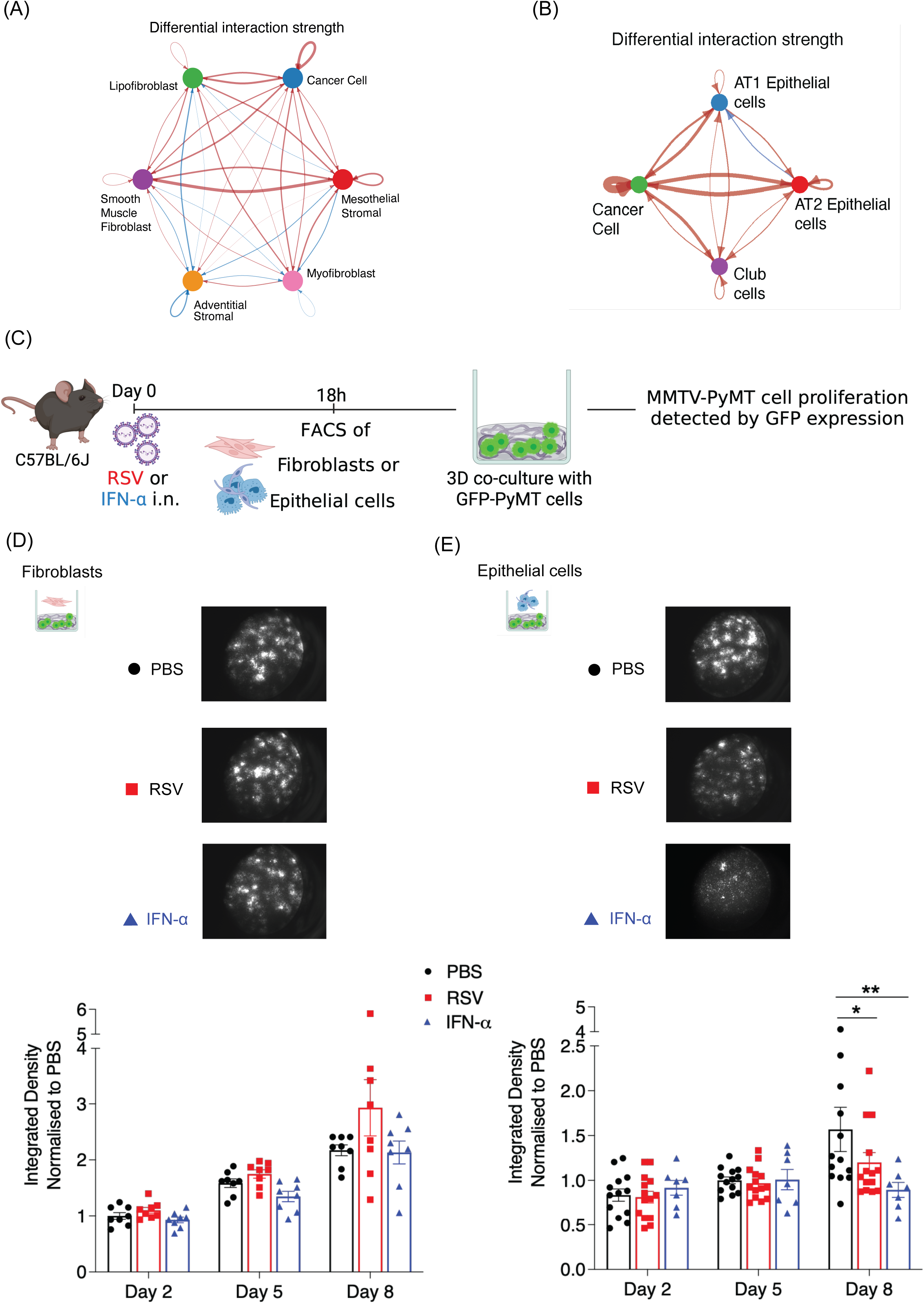
Lung epithelial cells from RSV infected mice or mice exposed to IFN-α are less supportive of tumor cell proliferation. C57BL/6J mice were intranasally treated with PBS (Control) or one dose of 1ug of IFN-α (IFN-α) and 18h later MMTV-PyMT cells were inoculated i.v. and 24h later the lungs cells were used for scRNA analysis. CellChat analysis was used to create Circle plot of Differential Interaction Score of Cancer Cells with stromal cell (A) or epithelial cell (B) compartments as based on Secreted ligands and Receptor interactions. (C) Experimental setup. IFN-α, PBS or RSV was intranasally administered to C57BL/6J. 18h later, fibroblasts and epithelial cells were sorted via FACS. Each cell type was co-cultured with GFP-expressing MMTV-PyMT cells in a collagen-solution-coated Alvatex Scaffold 96-well plate. Tumor cell proliferation was monitored daily for 8 days using the SteREO LumarV12 stereomicroscope. Images of fibroblasts (D) or epithelial cells (E) are representative of the co-cultures at day 8. All data points were normalized to the mean integrated density of the PBS group at day 2. Data are shown as mean +/- SEM. Data from fibroblast co-cultures are pooled from two independent experiments with 8 wells per condition. Data from co-cultures with epithelial cells are pooled from three independent experiments, n=13 for the PBS group, n=14 for RSV and n=7 for IFN-α. Statistical differences were assessed by two-way ANOVA with Tukey’s multiple comparisons. Only statistically significant differences are shown; *p<0.05; **p<0.01.

## Discussion

Under steady state, the lungs provide an environment that serves as an ideal niche for cancer metastases(2). However, the lungs can also be exposed to respiratory viruses, which cause acute inflammation and changes to the environment. The interplay between respiratory viral infections and the initiation and progression of lung metastasis remains unclear. Here, using the MMTV-PyMT experimental metastasis model, we show that early events during RSV infection led to reduced seeding and colonization of breast cancer cells in the lungs.

Viral infections induce early secretion of IFNs, especially type I IFNs(13). During RSV infection, AMs are the main producers of type I IFNs(17). As the type I IFN receptor is expressed on all nucleated cells, type I IFNs can have a global effect in all different lung cell types. Signaling through the type I IFN receptor potentiates type I IFN production through a positive feedback loop and induces expression of ISGs, which can limit viral replication. It also causes activation of neutrophils, inflammatory monocytes, NK cells, dendritic cells and macrophages, increasing the complexity of the lung environment(12,17,24,39). In this study we found that intranasal delivery of IFN-α could mimic the anti-metastatic effect induced by RSV infection in the lung, suggesting that type I IFN receptor signaling during RSV infection creates an unfavorable environment for cell metastases in this model. Interestingly, in several other metastatic tumor models (melanoma; B16, K1735m and DX3, fibrosarcoma; UV2237) it has been reported that prophylactic i.p. administration of human hybrid IFN-α also reduces the number of lung metastases, further supporting a role for type I IFNs in decreasing the seeding and colonization of metastatic cells(40–42). Furthermore, in hepatocellular carcinoma, subcutaneous treatment with IFN-α was shown to change the lung environment reducing the number of metastatic nodules independent of the primary tumor(26). There is evidence that type I IFNs can signal through the type I IFN receptor expressed on tumor cells to induce senescence or apoptosis(19). However, as intranasal administration of IFN-α to Ifnar1^-/-^ mice, (where only the MMTV-PyMT cells express the type I IFN receptor) had no impact on metastatic burden, the effect triggered by type I IFNs in our model is not due to a direct effect on tumor cells. Rather, type I IFNs mediate their anti-tumoral effect by changing the lung environment, leading to impaired establishment of metastatic breast cancer cell nodules.

We attempted to uncover the anti-tumoral role of the immune cells recruited and activated by RSV infection by depleting different immune cell populations. Interestingly, lack of RSV-induced neutrophil recruitment or depletion of neutrophils, NK cells, monocytes or T cells independently did not revert the phenotype and resulted in reduced numbers of metastatic foci. These depletion experiments demonstrate that none of the discussed immune cells on their own are necessary for mediating the anti-metastatic effect of type I IFNs and/or RSV infection. However, it is possible that due to the complexity of the lung environment during RSV infection, multiple immune cell types work together, or in a redundant manner, to impair metastatic initiation(43–45) and by reducing or depleting only one cell type, no overall effect on metastatic burden can be detected.

By measuring luciferase-expressing MMTV-PyMT cells *in vivo*, we demonstrated that exposure to RSV or IFN-α impairs tumor cell seeding and early proliferation in the lungs. Infiltrating cancer cells interact with cells in the lung parenchyma, which supports initiation of metastatic growth(46). For example lung epithelial cells play an essential role in supporting tumor cell growth(35,36,47) and they are also host cells for RSV replication(48). During respiratory viral infections, communication between lung epithelial cells, stromal cells and immune cells is critical for an effective viral clearance without excessive inflammation(38). We show that RSV- or IFN-α-primed lung epithelial cells are less supportive of MMTV-PyMT cell growth. The key changes in epithelial cells induced by how RSV infection or type I IFNs remain to be determined. This could be directly by signaling through the type I IFN receptor in epithelial cells or indirectly via type I IFNs activating other stromal cells, such as fibroblasts(49), which then act on epithelial cells. Nevertheless, altogether, our data demonstrates that impaired metastatic cell homing, survival and/or early proliferation in the lungs of RSV infected mice is likely mediated by rapid changes to the non-immune compartment induced by type I IFNs.

The urgent need to understand the interplay between respiratory viral infection and metastasis to the lungs is becoming increasingly evident after the COVID-19 pandemic.

Cohorts of cancer patients exposed to coronavirus infection provide a unique opportunity to follow cancer progression after respiratory viral infections. Notably, it has been proposed that COVID-19 disease can induce cancer reawakening and metastatic relapse(50–53). These human studies provide important insights and support our findings that respiratory viral infections can alter metastatic tumor progression. However, it is important to consider how different stages of cancer progression can dynamically alter the lung environment, perhaps leading to respiratory virus infections having differential effects that depend on the time point of infection relative to the stage of cancer. This is a key consideration when interpreting the data presented in our study, given that the MMTV-PyMT model lacks the generation of a pre-metastatic niche in the lungs by a primary tumor. However, this model has strengths in allowing us to directly investigate the temporal aspects of how respiratory viral infections impact on metastatic initiation, something that cannot be easily done in spontaneous metastasis models. Our findings may help identify targets that can be used to render the lungs less receptive to metastatic cells. Future studies will be necessary to assess how our findings can be translated into patients in clinical practice.

## Materials and methods

### Mice

For *in vivo* experiments performed at Imperial College London C57BL/6J, FVB/N and BALB/c mice were purchased from Charles River. Ifnar1^−/−^ mice (on a C57BL/6 background) and Myd88Trif^−/−^ mice (obtained from S. Akira (World Premier International Immunology Frontier Research Center, Osaka University, Osaka, Japan))(17) were bred in-house. For experiments held at The Francis Crick Institute, MMTV-PyMT mice on an FVB/N or C57BL/6J background, MMTV-PyMT actin-GFP on FVB/N background and MMTV-PyMT actin- luciferase also on the FVB/N background were bred in-house. The Myd88/Trif^-/-^ mice were Ifna6^gfp/+^ but since Ifna6 expression was not a primary readout the mice were designated as Myd88/Trif^-/-^ mice and compared to Ifna6^gfp/+^ wildtype (WT) mice. MMTV-PyMT mice on an FVB/N or C57BL/6J, bred at the Francis Crick Institute were obtained from The Jackson Laboratory(17). The MMTV-PyMT actin-GFP mice express the green fluorescent protein under the control of the actin promoter were originally a gift from J. Huelsken laboratory (EPFL, Lausanne, Switzerland) and the MMTV-PyMT actin-luciferase mice express firefly luciferase under the control of the actin promoter was a gift from D. Bonnet. For all experiments age-matched (7-12 weeks) female animals were used. All animal experiments were reviewed and approved by the Animal Welfare and Ethical Review Board (AWERB) within Imperial College London and approved by the UK Home Office in accordance with the Animals (Scientific Procedures) Act 1986 and the ARRIVE guidelines.

### Infections and treatments

Plaque-purified human RSV (originally A2 strain from ATCC, US) was grown in HEp2 cells as previously described(17). UV inactivated RSV (UV-RSV) was obtained by exposing the virus to UV light for 21min (UV RSV) in a CX-2000 UV cross-linker. For intranasal administration, FVB/N or C57Bl/6J mice were lightly anaesthetized and dosed with 6-7 x10^5^ focus-forming units (FFU) RSV or UV-RSV in 100 μl. For IFN-α (Miltenyi Biotec) exposure, one dose of 11µg or two doses of 500 ng 18h apart in 100 μl were intranasally administered(12).

### Antibody mediated neutrophil depletion

FVB/N mice were intraperitoneally (i.p.) treated with 150 µg in 100µl of anti-Ly6G (clone 1A8, Assay Genie, IE) or IgG isotype control (clone Y13/238, Cell services, the Francis Crick Institute) a day before RSV infection and every second day until day 4 post infection.

### Antibody mediated monocyte depletion

C57BL/6J mice were treated with 20 µg of anti-CCR2 (clone MC21(29)) or isotype-matched control rat IgG2b (Assay Genie, IE) in 100µl i.p. 6h before RSV infection and daily until day 5 p.i..

### Antibody mediated NK cell depletion

C57BL/6J mice were treated with 100 µg of anti-NK1.1 (clone PK136, eBioscience) or isotype mouse IgG2a (clone C1.18.4, BioXcell) in 100µl i.n. 24h before RSV infection. Mice were treated with 200 µg in 200 µl of anti-NK1.1 or isotype control i.p. the day before infection and daily until day 4 post infection.

### Antibody mediated T cell depletion

C57BL/6J mice were exposed to PBS or RSV i.n.. Depletion of T cells was performed by i.p. administration of 150 µg of anti-CD4 (clone GK1.5, Assay Genie, IE) and 150 µg of anti-CD8 (clone YTS 169) from the day of infection until day 6 p.i. every alternate day. Isotype-matched control of 300 µg rat IgG2b (Assay Genie, IE) was performed following the same regime.

### Experimental lung metastases

MMTV–PyMT primary cells, isolated from MMTV–PyMT breast tumors as previously described(46) were used in all experiments. Briefly, tumors were mechanically digested with a razor blade and digested with DNase (37.5ml/ml, Roche Diagnostics), Liberase Tm (75ml/ml, Roche Diagnostics), and Liberase Th (75 ml/ml, Roche Diagnostics), filtered through a 100μm filter and washed twice in DMEM containing 10% fetal calf serum (FCS). Cells were cultured overnight on collagen coated dishes (30 μg/ml PureCol collagen (Advanced Biomatrix), 0.1% bovine serum albumin (BSA, Sigma) and 20 mM HEPES in HBSS (Thermo Fisher Scientific)) in MEM media (DMEM/F12 (Thermo Fisher Scientific) with 2% FCS, 100 U/ml penicillin–streptomycin (Thermo Fisher Scientific), 20 ng/ml EGF (Thermo Fisher Scientific) and 10 μg/ml insulin (Merck Sigma-Aldrich). The following day, cells were washed with PBS followed by a 5 min wash of 1 mM EDTA at 37°C. MMTV-PyMT cells were detached with 0.05% Trypsin-EDTA (Gibco) for 7 min at 37°C. Cells were washed with DMEM+10% FCS and frozen in 10% DMSO, 40% FCS, 50% MEM media.

4T1 and 4T1-GFP expressing cells were provided by the Cell Services Unit of The Francis Crick Institute. Cells were grown in DMEM media, supplemented with 10% FCS, 2 mM L- glutamine, 1001U/ml penicillin, and 100 μg/ml streptomycin. For experimental lung metastases, 0.1x10^6^ cells were intravenously injected in 200μl of PBS.

For tail vein injections, cells were thawed and cultured overnight on collagen-coated dishes as previously described. After detachment, cells were washed twice in PBS and filtered through a 100 μm filter (pluriSelect). For all experiments, except for *in vivo* imaging (see below), 0.3x10^6^ cells, in 200μl, were inoculated intravenously (i.v.). For experimental lung metastases using MMTV-PyMT-GFP expressing cells, FVB/N mice were i.v. injected with 1x10^6^ cells in 200μl.

### *in vivo* luminescence imaging

These experiments were performed at the Francis Crick Institute. Mice were anaesthetized with IsoFlo (isoflurane, Abbott Animal Health) and intranasally infected with 4x10^5^ FFU of RSV or treated with 1 µg of IFN-α in 50 µl and 24h later 0.5x10^6^ MMTV-PyMT actin- luciferase cells were inoculated i.v.. To measure luciferase activity, at different times post tumor cells injection, animals were administered 100 ul of 30 mg/ml D-Luciferin (Xenogen) i.p.. After 12 minutes the animals were anaesthetized with inhaled isoflurane. The anaesthetized animals were then imaged using IVIS Spectrum (Perkin Elmer)(54). At different times post tumor cells injection luminescence signal (photons/second) was quantified using Living Image Software (Perkin Elmer) by selecting a ROI of a set size positioned over the thorax of each animal. *ex vivo* bioluminescence intensity was measured to assess metastatic burden at 28 days post cell administration(54).

### Metastases burden analysis

Mice were sacrificed 28 days after tumor cell injection by a fatal dose of pentobarbital injected i.p.. Lungs were inflated with 1.5ml of PBS, fixed in 10% Formalin (Sigma-Aldrich) for 16h and embedded in paraffin blocks. Three sections of 4μm at least 150μm apart were stained for H&E according to standard procedures. Sections were scanned using the Axio Scan Z1 slide scanner (Zeiss, Germany).

Macroscopic metastatic burden was quantified by counting nodules on the surface of the lung using the Zeiss SteREO Lumar.V12 stereoscope and is shown as metastatic nodules per lung per mouse. For number of microscopic metastatic nodules and size distribution, three H&E-stained sections of 4 μm at least 150 μm apart were analyzed using the Zen Blue software (Zeiss, Germany). Metastatic tumor burden is shown as fold change to the average of metastatic nodules in the PBS group in each experiment.

### Isolation of lung immune cells and airway cells

For flow cytometry, lungs were perfused with PBS and collected in complete DMEM (cDMEM; supplemented with 10% FCS, 2 mM L-glutamine, 1001U/ml penicillin, and 100 μg/ml streptomycin). Lungs were processed with a gentle MACS dissociator (Miltenyi Biotech) according to manufacturer’s protocol and digested with Collagenase D (11mg/ml; Roche) and DNase I (30 µg/ml; Invitrogen) for 11h shaking at 37°C and processed again in a gentle MACS dissociator. Red blood cells were lysed using an ACK buffer (0.15 M NH_4_Cl, 1.0 mM KHCO_3_, 0.11mM Na_2_EDTA). Cells were re-suspended in PBS and filtered through a 1001µM cell strainer (Greiner BioOne) and the absolute number of recovered cells was quantified by Trypan Blue (Thermo Fisher Scientific) exclusion of dead cells.

Bronchoalveolar lavage (BAL) was performed by flushing 1 ml of PBS with 0.5 mM EDTA through the trachea 3 times. The BAL fluid was centrifuged, the supernatant was stored at −80 °C and the cells were treated with ACK buffer to remove red blood cells. Total airway cell number was determined by Trypan Blue (Thermo Fisher Scientific) exclusion of dead cells. Lung cells and airway cells were further analyzed by flow cytometry.

### Flow cytometry

For immune cell characterization, 2.5×10^6^ lung cells or all recovered BAL cells were incubated with a purified rat IgG2b anti–mouse CD16/CD32 receptor antibody (BD) for 20 min at 4°C in FACS buffer (PBS supplemented with 1% BSA and 0.05 mM EDTA). For surface staining, cells were stained in PBS with fixable live-dead Aqua dye (Invitrogen) and with different fluorochrome-conjugated antibodies (see Supp. Table 1) for 25 min at 4°C. Following staining, cells were fixed with 100 µl 1% paraformaldehyde or Cytofix™ Fixation Buffer (BD) for 201mins at 4°C and stored in FACS buffer. Analysis was performed on a BD LSR Fortessa, by acquiring 2.5x10^5^ single, live CD45^+^ cells. Data were analyzed with FlowJo software (Tree Star). Total cell populations were quantified as the whole lung count x (proportion of lung tissue sampled) x1(%CD45^+^ cell of live cells)1×1(% of population of interest out of CD45^+^ cells).

### Detection of tumor cells by flow cytometry

To quantify tumor cells, lungs were collected at the specified time points after tumor cell injection and digested with Liberase Tm (Roche Diagnostics), Liberase Th (Roche Diagnostics) and Deoxyribonuclease I (Merck Sigma-Aldrich). in HBSS solution for 30 min at 37°C with 180 rpm agitation. The digested mixture was then passed through a 100μm strainer, washed with 10% FCS HBSS solution and centrifuged for 8 min at 300 x g. Cells were then resuspended in Red Blood Cell Lysis Solution (Miltenyi Biotec) for 5 min at room temperature and then passed through a 40μm strainer. After centrifugation, cells were washed with MACS buffer (0.5% BSA and 2 mM EDTA in PBS) and passed through a 20μm strainer-capped tube to generate a single-cell suspension. All recovered cells were stained as described in the flow cytometry section but acquired immediately after staining without fixing.

### RNA extraction and quantitative RT-PCR

For RNA extractions, lung lobes were snap-frozen in liquid nitrogen and stored at −80°C. Lungs were homogenized using a TissueLyser LT (Qiagen) and total RNA was extracted from the lung tissue supernatant using RNeasy Mini kit (Qiagen) including a DNase digestion step. RNA yield was determined by NanoDrop (Thermo Scientific). Retro transcription to cDNA of 1 or 2 µg of RNA was performed using High Capacity RNA-to-cDNA kit (Applied Biosystems). Real time PCR was performed using Quantitect Probe PCR Master Mix (Qiagen) in the 7500 Fast Real-Time PCR System (Applied Biosystems). Detection of mRNA of Gapdh (encoding glyceraldehyde-3-phosphate dehydrogenase), Cxcl1, Ccl2, Il1b, Ifna5, Cxcl10, Mx1 and Il6 was achieved using specific primers and probes (all Applied Biosystems). Relative mRNA expression of the genes of interest are shown as 2^−ΔCT^. Quantitative PCRs for Ifnb, Ifnl, Oas1, Rsad2 (Viperin) and Eif2ak2 (Pkr) and RSV L gene was performed using primers and probes previously described(17). Analysis was performed using 7500 Fast System SDS Software (Applied Biosystems). Number of copies of the gene of interest are shown as copy number per ug of RNA and was calculated using a plasmid DNA standard curve for each gene and normalized to Gapdh, as previously described(12).

### Immune mediator detection

Levels of IFN-γ, Granzyme B (both DuoSet ELISA kits from R&D systems), IL-6(12) and IFN-α(24) present in the BAL fluid were measured by ELISA. Absorbance was determined at 450 nm, on FLUOstar Omega (BMG Labtech) plate readers and analyzed using Mars (BMG Labtech) software.

### 3D cell culture

MMTV-PyMT GFP^+^ cells were plated (5,000 cells per well) into a collagen-coated 96-well Alvetex Scaffold plate (ReproCELL)(36). On the following day, lungs were harvested 18h post PBS, IFN-α or RSV exposure and enzymatically digested, as described in the section ‘Detection of tumor cells by flow cytometry’. Cells were then stained with CD45 and CD31 MicroBeads (Miltenyi Biotec) for magnetic depletion of immune and endothelial cells according to manufacturer’s protocol and then stained with a fluorescent-conjugated EpCAM antibody for FACS sorting(6, 7). Purity after all sorts were above 95%. 50,000 lung fibroblasts (CD45^-^CD31^-^EpCAM^-^) or lung epithelial cells (CD45^-^CD31^-^EpCAM^+^) were added to the scaffolds. Results are normalized to the average GFP expression at day 2 of the PBS condition of each cell type for each independent experiment.

### scRNAseq

Liberase-digested lung cells pooled from 5 mice per group were stained and then fixed for 16hrs using a 4% formaldehyde fixative solution, as described in the Demonstrated Protocols CG000478 and CG000553 and using Chromium Next GEM Single Cell Fixed RNA Sample Preparation Kit (10x Genomics PN-1000414), prior to sorting (BD Influx). Cells were sorted based on CD45^+^ (leukocytes), EpCAM^+^ (epithelial), CD45^-^EpCAM^-^ (mesenchymal) and GFP (cancer cells) and added to the final sample at a ratio of 3:3:3:1. Cells were pooled form 3 independent experiments. Quench buffer (10x Genomics, PN-2000516) was added to the sample for storage at -80°C.

Gene expression was measured using barcoded probe pairs designed to hybridize to mRNA specifically. Using a microfluidic chip, the fixed and probe-hybridized single cell suspensions were partitioned into nanolitre-scale Gel Beads-in-emulsion (GEMs). A pool of ∼737,000 10x GEM Barcodes was sampled separately to index the contents of each partition. Inside the GEMs, probes were ligated and the 10x GEM Barcode was added, and all ligated probes within a GEM share a common 10x GEM Barcode. Barcoded and ligated probes were then pre-amplified in bulk, after which gene expression libraries were generated (User Guide: CG000527, Chromium Fixed RNA Profiling Reagent Kits for Multiplexed Samples) and sequenced (NovaSeq 6000. Sequencing read configuration: 28-10-10-90).

### Bioinformatic Analysis

Raw reads were initially processed by the Cell Ranger v.2.1.1 pipeline, which deconvolved reads to their cell of origin using the UMI tags, aligned these to the mm10 transcriptome using STAR (v.2.5.1b) and reported cell-specific gene expression count estimates. All subsequent analyses were performed in R v.4.3.3 using the cellrangerRkit packages. Genes were ‘expressed’ if the estimated (log_10_) count was at least 0.1. Primary filtering was then performed by removing from consideration: genes expressed in fewer than 3 cells; cells expressing fewer than 200 genes; cells for which the total yield (that is, sum of expression across all genes) was more than two standard deviations from the mean across all cells in that sample; and cells for which mitochondrial genes made up greater than 4% of all expressed genes. Seurat V5(55) was used to perform UMAP and for nearest-neighbour analysis on the integrated datasets (rPCA). Clusters identified using resolution of 0.5 were either assigned manually using specific cell type expression or for immune cells by cross-referencing to the ImmGen dataset(56). fGsea package was used for identification of cancer cell pathway changes using gene expression changes from FindMarkers and cross-referencing to Reactome pathways(57). CellChat (58) was used to identify receptor-ligand interaction changes between the two conditions.

### Statistical analysis

Statistical analysis was performed using Prism (GraphPad software) version 9. Data are presented as the mean1+/-1SEM. As indicated in each figure legend, two tail unpaired Student’s t-test, a two-way ANOVA with mixed-effect analysis or a One-way ANOVA was performed. Only p values1<10.05, considered statistically significant for all tests are shown in the figures.

## Author contribution

A.F. designed, performed and analyzed the experiments and wrote the paper. V.B. and F.R., designed, performed and analyzed specific experiments and reviewed the paper. A.O., S.R. and R.F. performed specific experiments and reviewed the paper. M.M. provided the anti- CCR2 depletion antibody I.M. designed the experiments and reviewed the paper. C.J. supervised the project, designed the experiments, and wrote the paper.

## Acknowledgements

We thank S. Akira (World Premier International Immunology Frontier Research Center, Osaka University, Osaka, Japan) for providing Myd88/Trif^-/-^ mice. We thank the St Mary’s flow cytometry facility and the staff of the St Mary’s animal facility for their assistance as well as the Experimental Histopathology lab and the BRF of the Francis Crick institute for their support.

We thank Christina Michalaki, Minerva Garcia Martin and Sophie Guan for help with experiments and Franz Puttur and Caetano Reis e Sousa for critically reading the manuscript. We thank members of the Johansson and Malanchi lab for scientific discussions. Biorender was used for depicting the experimental setups.

