## Supplemental information for "Type I interferons induced upon respiratory viral infection impair lung metastatic initiation"

**Supplementary Material**

**Supp. Table 1. Antibodies used for flow cytometry**, for gating strategies refer to Supp. Fig. 2, 3, 5 ,6, 10 and 12.

| Target | Clone | Fluorochrome | Company | Working concentration |
|---|---|---|---|---|
| CD45 | 30-F11 | BV605 | Biolegend | 0.25 µg/ml |
| CD45 | 30-F11 | PerCP-Cy5.5 | Biolegend | 0.25 µg/ml |
| Syglec F | E50-2440 | BV786 | BD Bioscience | 0.5 µg/ml |
| Ly6G | 1A8 | Alexa Fluor 488 | Biolegend | 5 µg/ml |
| CD64 | X54-5/7.1 | APC | Biolegend | 1 µg/ml |
| CD64 | X54-5/7.1 | PE | Biolegend | 1 µg/ml |
| CD11c | HL3 | V450 | BD Bioscience | 1 µg/ml |
| CD11b | M1/70 | PE-Cy7 | eBioscience | 0.5 µg/ml |
| CD3 | 145-2C11 | APC-efluor780 | eBioscience | 2 µg/ml |
| CD19 | 1D3 | APC | Biolegend | 1 µg/ml |
| CD3 | 17A2 | Alexa Fluor 700 | eBioscience | 2 µg/ml |
| CD8 | 53-6.7 | eFluor780 | eBioscience | 0.5 µg/ml |
| CD4 | GK1.5 | PE | eBioscience | 0.5 µg/ml |
| CD4 | RM4-5 | APC | BD Bioscience | 0.5 µg/ml |
| CD69 | H1.2F3 | BUV737 | BD Bioscience | 1 µg/ml |
| CD49b | DX5 | PE | Biolegend | 1 µg/ml |
| Ly6C | 12HK1.4 | BV711 | eBioscience | 0.5 µg/ml |
| EpCAM | G8.8 | APCFire750 | eBioscience | 2 µg/ml |
| EpCAM | G8.8 | APC | eBioscience | 2 µg/ml |
| PD-1 | BV605 | 29F.1A12 | Biolegend | 2 µg/ml |

**Supplementary data**

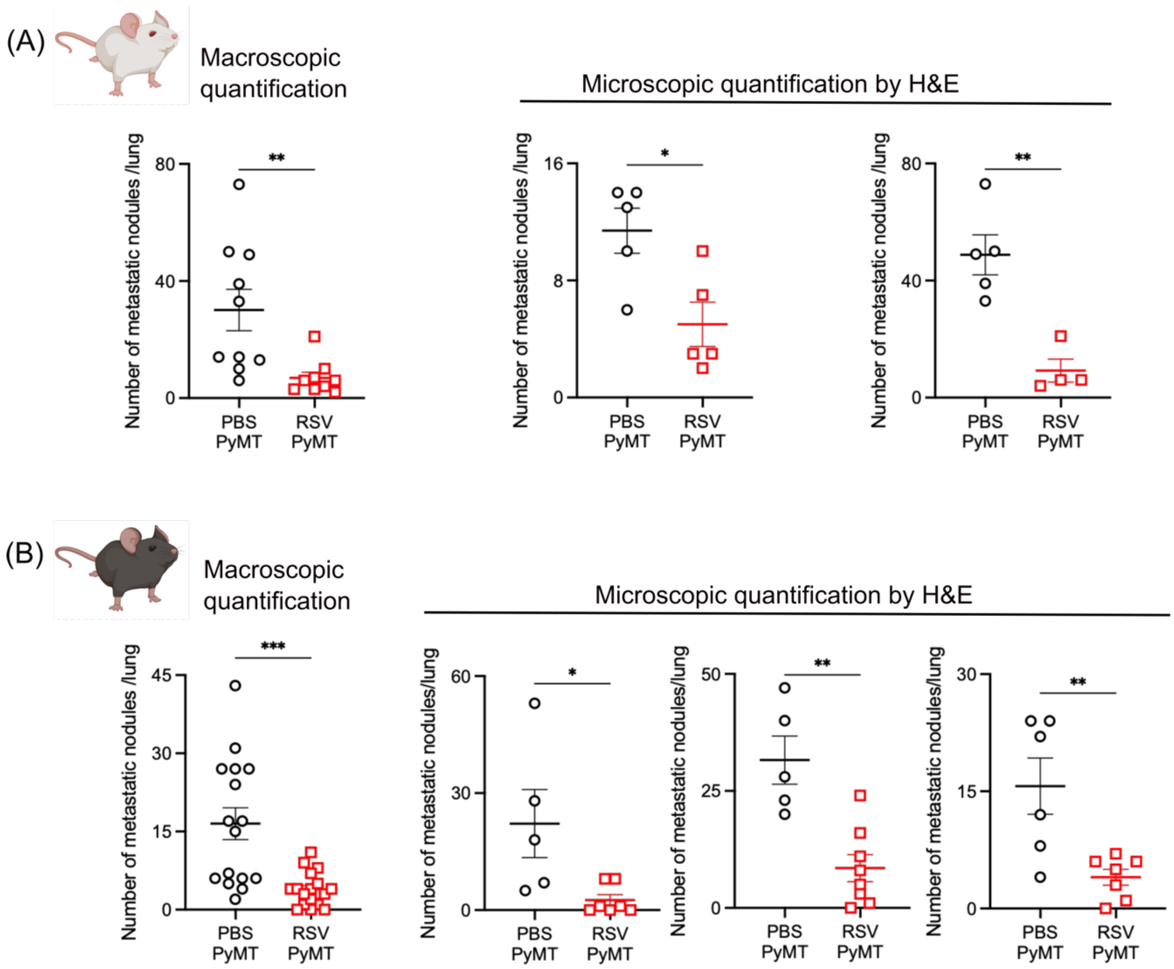


**Supp. Fig. 1. Macroscopic and microscopic quantification of metastatic MMTV-PyMT nodules in mock or RSV infected mice.** Lungs were harvested from PBS (PBS PyMT), or RSV (RSV PyMT) exposed i.n. (A) FVB/N and (B) C57BL/6J mice that received MMTV-PyMT cells i.v. a day later. Gross macroscopic quantification of lung metastatic nodules was performed. Also, microscopic quantification performed by quantifying nodules in H&E-stained lung sections is shown for each independent experiment. These data were normalized and pooled in Fig. 1. Data shown for macroscopic count are pool from two independent experiments n=10 for PBS and 9 RSV infected FVB/N mice and from three independent experiments n=16 for PBS and 23 for RSV infected C57BL/6J mice. All data are shown as mean +/-SEM. Student's t-test statistical analysis was performed; *p<0.05, **p<0.01, ***p<0.0005.

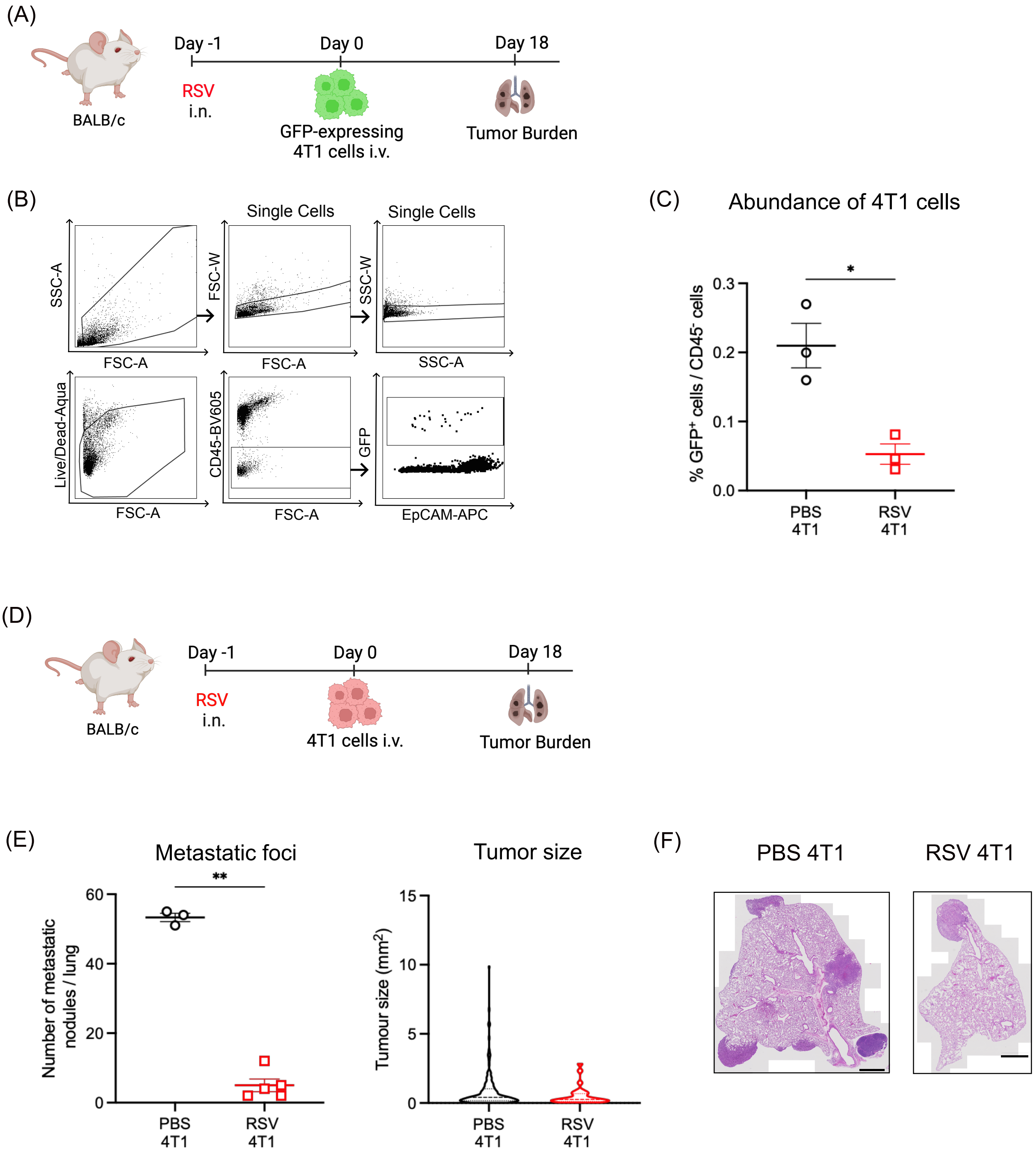


**Supp. Fig. 2. RSV infection impairs 4T1 experimental lung metastasis.** (A) BALB/c mice were infected i.n. with RSV or mock infected (PBS). A day later mice were injected i.v. with $1x10^5$ GFP expressing 4T1 cells. Abundance of tumor cells in the lungs was assessed a week later by flow cytometry. (B) Gating strategy and (C) percentage of $GFP^+$ 4T1 cells in the lungs of infected or control mice. (D) The same set up was used with unlabelled 4T1 cells. (E) Number of metastatic nodules and their size was assessed by histological analysis after 18 days. (F) Representative H&E-stained section of the lung, scale bar 1000mm. Data from one experiment shown as mean +/- SEM, with (C) 3 mice per group and (E) 3 mice in the control group and 5 mice in the infected group. Student's *t*-test was performed. Only statistically significant differences are shown; *p<0.05, **p<0.01

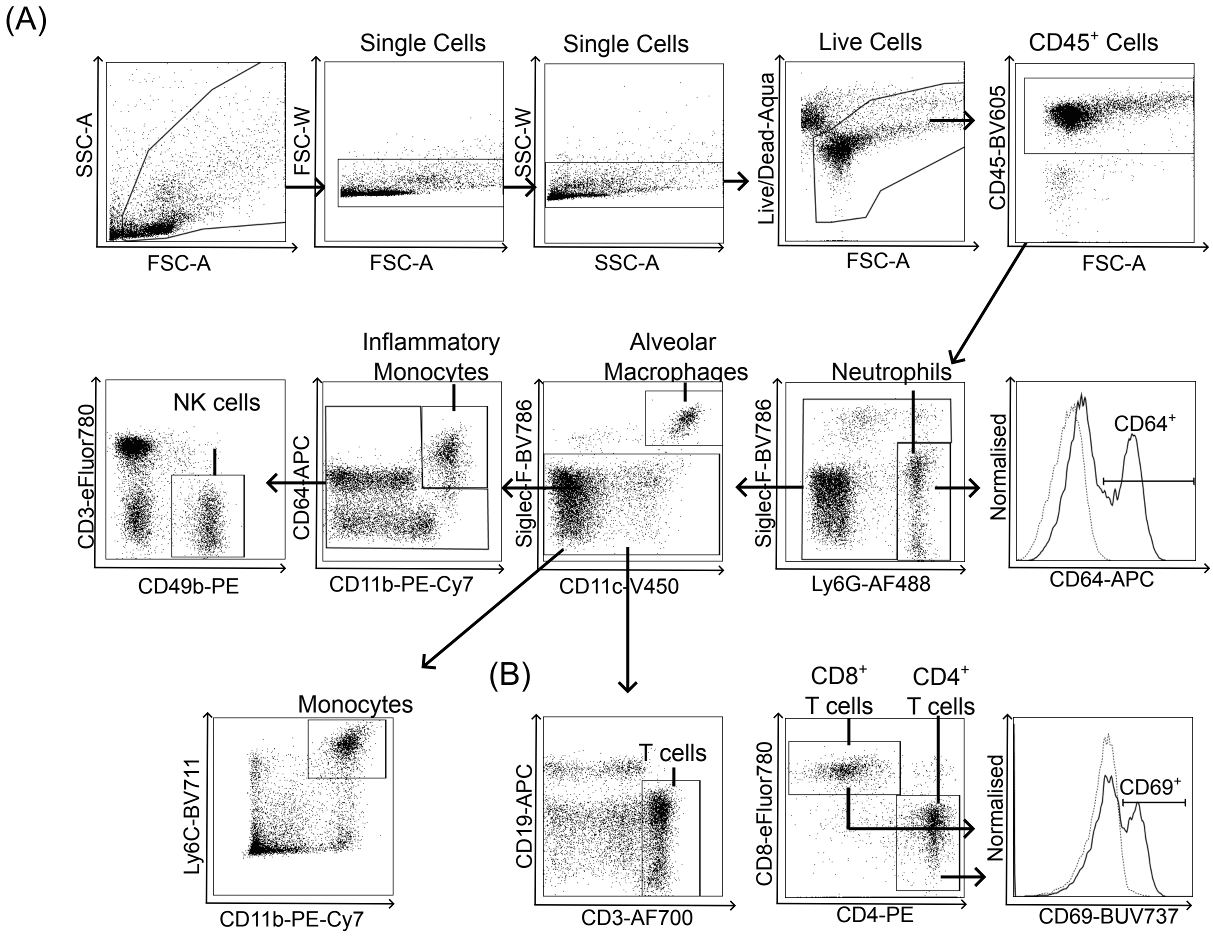


**Supp. Fig. 3. Gating strategy to identify cell populations in lungs and airway during RSV infection in the presence of MMTV-PyMT cells.** Shown are representative flow cytometry plots from lung cells harvested (A) 18h or (B) 8 days post RSV infection. (A) Flow cytometry analysis depicting the frequency of live CD45$^+$ cells after excluding debris, doublets and dead cells and downstream gating strategy used to define neutrophils, activated neutrophils as CD64$^+$, alveolar macrophages (AMs), inflammatory monocytes, gated as Ly6C$^+$ CD11b$^+$ or CD11b$^+$ CD64$^+$ and NK cells. (B) Gating strategy showing proportion of CD8$^+$ and CD4$^+$ T cells and histogram plot showing CD69 expressing CD8$^+$ T cells, the same analysis was performed for CD4$^+$ T cells. Data shown from an RSV infected mouse 8 days post infection.

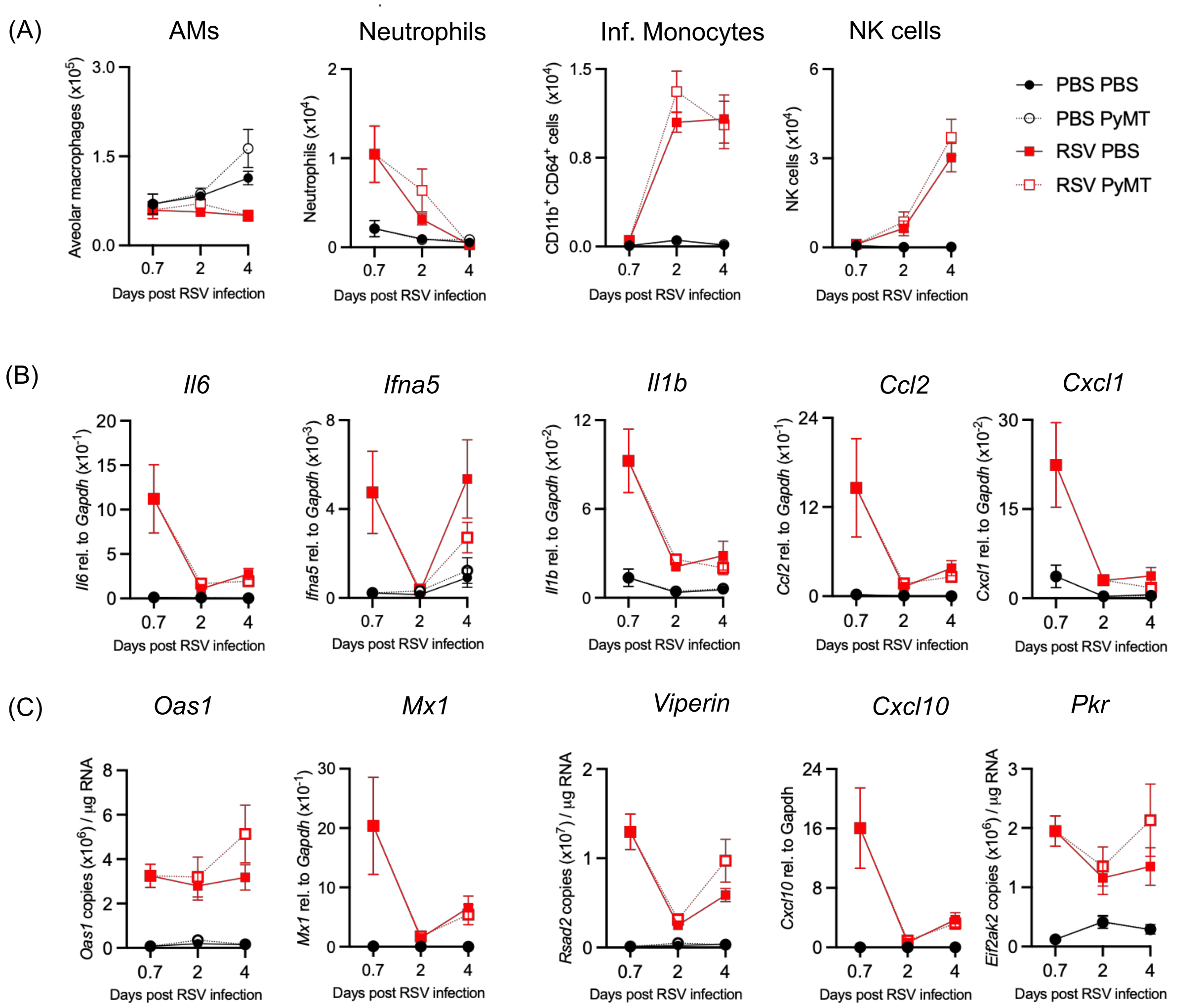


**Supp. Fig. 4. Immune response in lungs and airways during RSV infection in the presence of MMTV-PyMT cells.** FVB/N mice were infected i.n. with RSV or mock infected (PBS). A day later 3x10$^5$ MMTV-PyMT cells (PyMT) or PBS were inoculated i.v.. (A) Cells present in the airways were recovered from the BAL, at 0.7, 2, 4 and 8 days p.i. and stained for surface markers to enumerate different airway leukocyte populations, as shown in Supp. Fig. 3A. Shown are number of alveolar macrophages (AMs), neutrophils, inflammatory (Inf) monocytes and NK cells at different times post RSV infection. (B) Expression of *Il6, Ifna5, Il1b, Ccl2,* and *Cxcl1 and* (C) the interferon stimulated genes *Oas1, Mx1, Viperin, Cxcl10* and *Pkr*, were quantified in RNA from lungs by RT-qPCR. Data for day 0.7 are pooled from two independent experiments presented as the mean +/- SEM of 9 mice per group. Data for days 2 and 4 are pooled from two independent experiments with 8 mice per group. One-way ANOVA was performed to compare the infected groups followed by Tukey's post hoc test at each time point, no differences were detected.

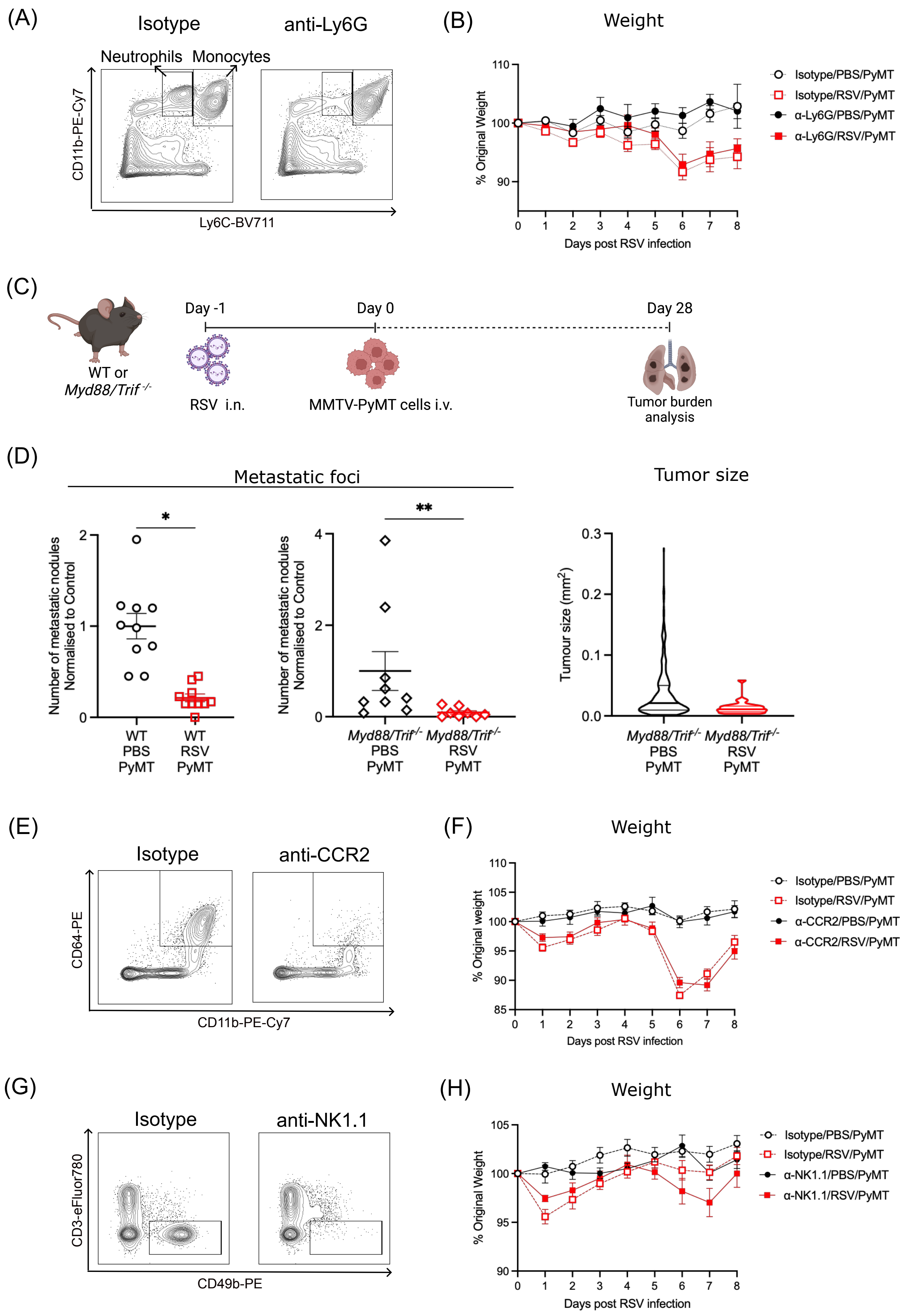


**Supp. Fig. 5. Lower number of metastatic nodules is not due to RSV induced recruitment of neutrophils, monocytes or NK cells.** (A) FVB/N mice were treated i.p. with anti-Ly6G or isotype control every second day from the day prior to infection. Representative flow cytometry of CD11b$^+$ Ly6C$^+$ cells gated from lungs live cells, CD45$^+$, SiglecF$^-$ on day 2 p.i. (B) Weight loss of mice treated with anti-Ly6G or isotype control, infected with RSV followed by i.v. administration of tumor cells a day later. (C) *Myd88/Trif*$^{-/-}$ or wildtype (WT) mice were exposed i.n. with PBS or RSV at day -1 and then inoculated with MMTV-PyMT cells (PyMT) i.v. on day 0. Lungs were analyzed for tumor burden after 28 days. (D) Number of metastatic nodules and tumor size were quantified 28 days post tumor inoculation by H&E staining. Student's t-test statistical analysis was performed *p<0.05, **p<0.01. (E) Monocytes were depleted in C57BL/6J mice using anti-CCR2 from the day of infection. Representative dot plot from CD11b$^+$ CD64$^+$ cells in lungs, gated from lungs live cells, CD45$^+$, SiglecF$^-$, Ly6G$^-$ at day 2 p.i. (F) RSV infection was followed by weight loss until day 8 p.i.. (G) C57BL/6J mice were depleted of NK cells following i.p. and i.n. the day prior to infection, and then i.p. administration of anti-NK1.1 antibody daily from day -1 to day 3. Mice were infected with RSV or mock-infected (PBS) at day -1 and inoculated with tumor cells at day 0. Depletion of NK cells in the lung was analyzed by flow cytometry at day 4 p.i., gated from lungs live cells, CD45$^+$, SiglecF$^-$, Ly6G$^-$, CD3$^-$, CD49b$^+$. (H) Disease severity after RSV infection was followed by weight loss. Data in (D) are pooled from two independent experiments and shown as mean +/-SEM of n=10 in each wildtype group and n=9 for the *Myd88/Trif*$^{-/-}$PBS group and n=8 for the *Myd88/Trif*$^{-/-}$ RSV group. In (B, F, H) a two-way ANOVA, mixed-effect analysis was performed to compare weight loss after infection followed by Tukey's post hoc test, with no significant differences detected. For n numbers refer to Fig. 4.

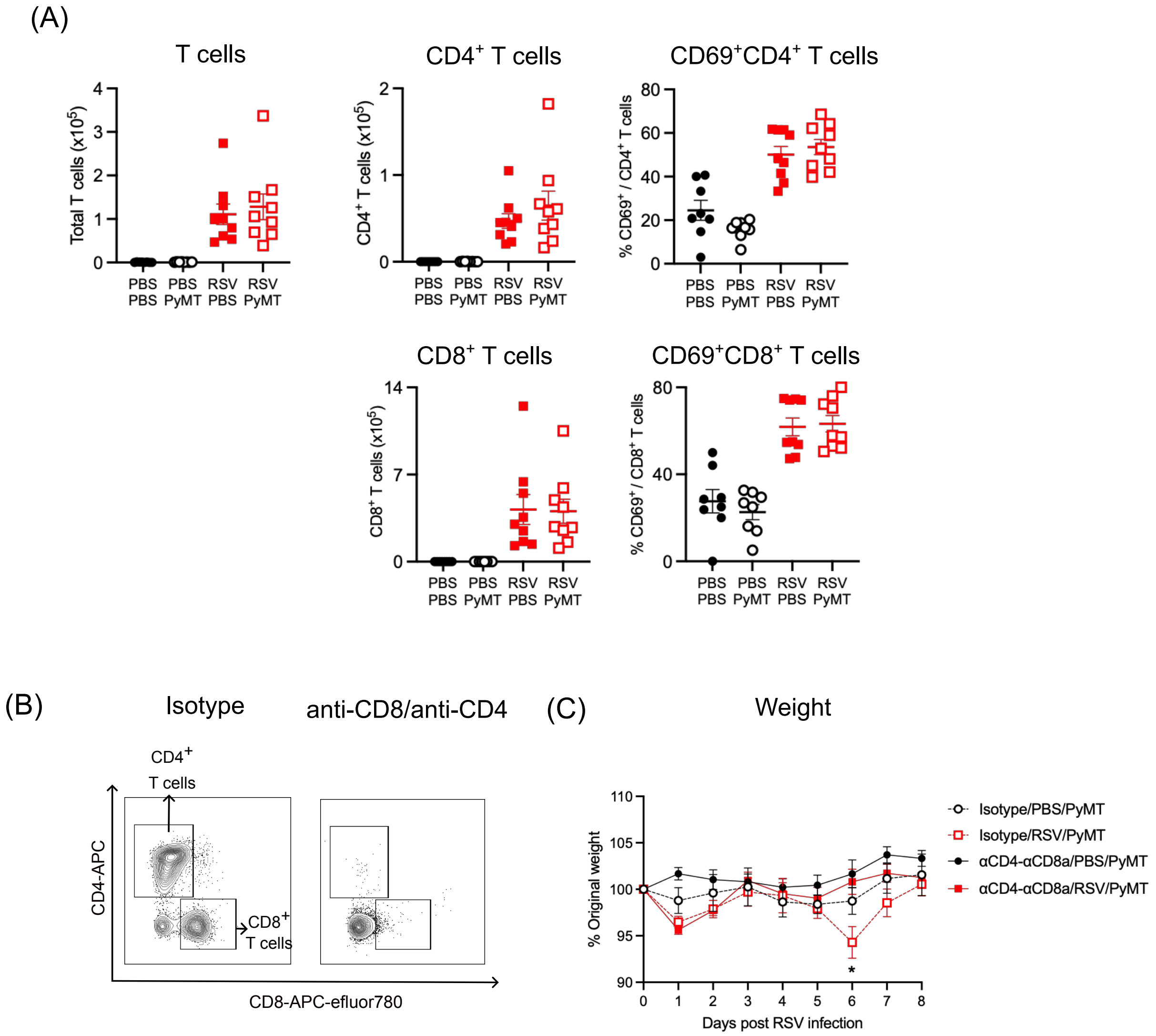


**Supp. Fig. 6. T cell responses in the airways during RSV infection in the presence of MMTV-PyMT cells.** FVB/N mice were infected i.n. with RSV or mock-infected (PBS). A day later MMTV-PyMT cells or PBS were inoculated i.v.. (A) Number of total T cells, CD4$^+$ T cells, CD69$^+$CD4$^+$ T cells, CD8$^+$ T cells and CD69$^+$CD8$^+$ T cells were quantified in BAL 8 days p.i., following the gating strategy shown in Supp. Fig. 3B. Data are pooled from two independent experiments, with n=8 for both uninfected group and n=9 for both RSV infected mice. Student's t-test was performed to compare the two RSV infected groups, with no differences detected. (B) CD4$^+$ and CD8$^+$ T cells were depleted during RSV infection using the respective antibodies or isotype control every second day, from the day of infection until day 6 p.i. Representative dot plot showing CD4$^+$ and CD8$^+$ T cells from lungs at day 7 p.i., gated from live, CD45$^+$, SiglecF$^-$, Ly6G$^-$, CD3$^+$. (C) Weight loss after RSV infection was followed daily until day 8 p.i.. Data are representative from two independent experiments presented as the mean +/- SEM of 9 mice per group in the PBS group and 8 or 10 mice in RSV infected mice treated with isotype or anti-CD4 and anti-CD8, respectively. A two-way ANOVA, mixed-effect analysis was performed to compare weight loss after infection in all groups, followed by Tukey's post hoc test, *p<0.05.

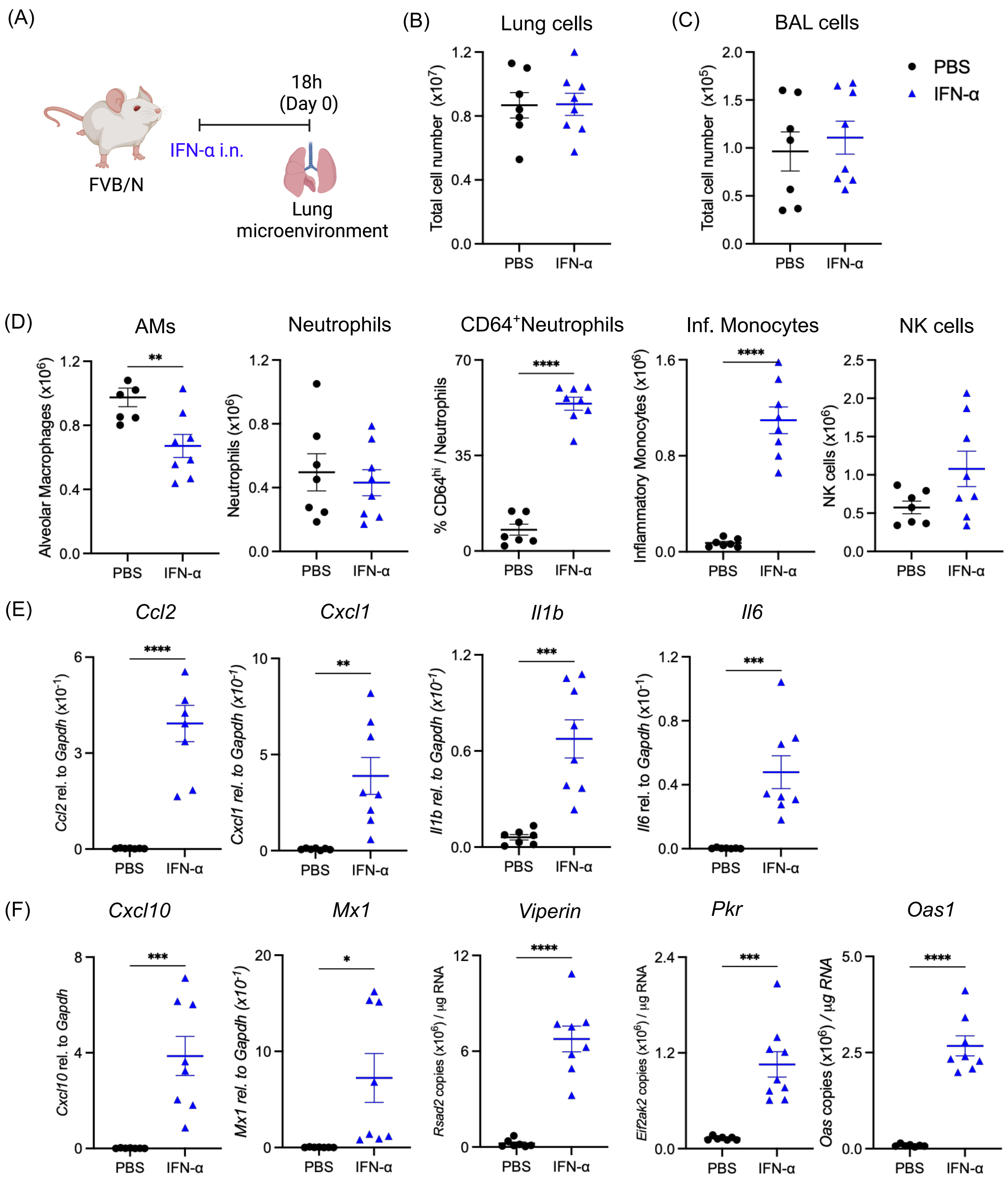


**Supp. Fig. 7. Recombinant IFN-α administered i.n. induce transient changes in the lung microenvironment.** FVB/N mice were treated i.n. with 1ug of IFN-α and 18h post treatment lungs and BAL were obtained. (A) Schematic representation of the treatment regime. At 18h, total number of (B) lung cells and (C) BAL cells were quantified. (D) AMs, neutrophils, CD64+ neutrophils, inflammatory (Inf) monocytes and NK cells were quantified after PBS or IFN-α exposure. (E) Relative gene expression of *Ccl2, Cxcl1, Il1b, Il6* and (F) relative gene expression of gene copy number of the interferon stimulated genes *Mx1, Viperin, Pkr, Cxcl10* and *Oas1* were quantified and normalized to housekeeping gene *Gapdh* or quantified using a standard curve. PBS data are also shown as PBS day 0.7 in Fig. 2. Data shown are pooled from two independent experiments with n=6 for PBS and n=7 for IFN-$\alpha$ +/-SEM. Student's *t*-test analysis was performed. Only statistically significant differences are shown; *p<0.05, **p<0.01, ***p<0.005, ****p<0.0001.

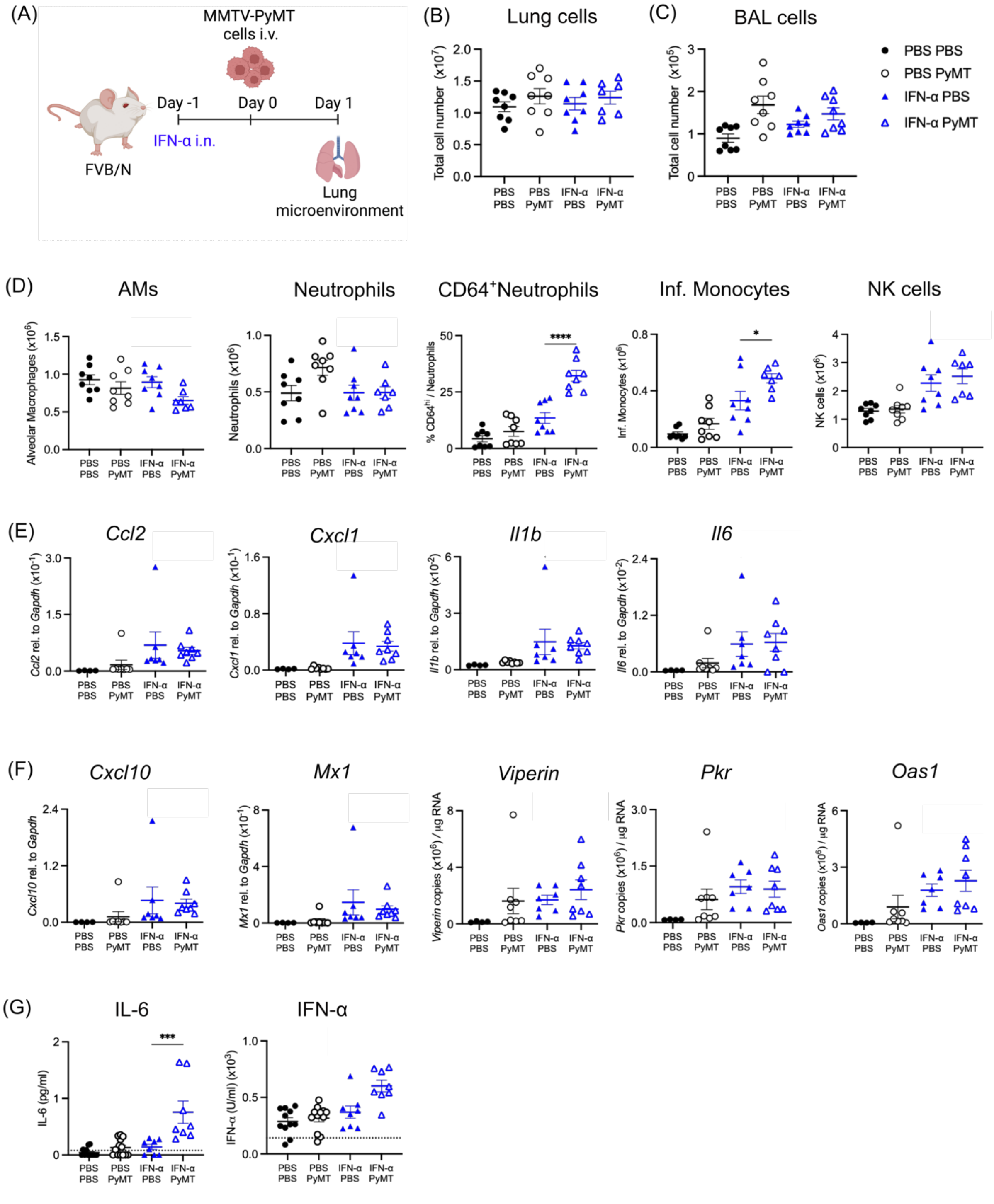


**Supp. Fig. 8. Presence of MMTV-PyMT cells in the lungs do not change the response induced by IFN-α** FVB/N mice were treated i.n. with 1ug of IFN-α at day -1, inoculated with tumor cells at day 0 and lungs and BAL were harvested 18 hours later. (A) Schematic representation of the treatment regime. At 18h post MMTV-PyMT cell administration, total number of (B) lung cells and (C) BAL cells were quantified. In (D) innate immune cell in the lungs were quantified by flow cytometry. (E) Relative gene expression of *Ccl2, Cxcl1, Il1b, Il6* and (F) relative gene expression or gene copy number of the interferon stimulated genes *Cxcl10, Mx1, Viperin, Pkr and Oas1* were quantified and normalized to housekeeping gene *Gapdh*. Number of copies were calculated using a plasmid standard curve for *Viperin*, *Pkr* and *Oas1*. (G) Concentration of IL-6 and IFN-$\alpha$ was quantified in BAL fluid, dotted line shows detection limit. PBS data are also shown as PBS day 2 in Fig. 2. Data shown are representative from two independent experiments with n=8 +/-SEM. One-way ANOVA with Tukey's post hoc test was performed to compare the IFN-$\alpha$ exposed groups. Only statistically significant differences are shown; * $p<0.05$; *** $p<0.005$; **** $p<0.0001$.

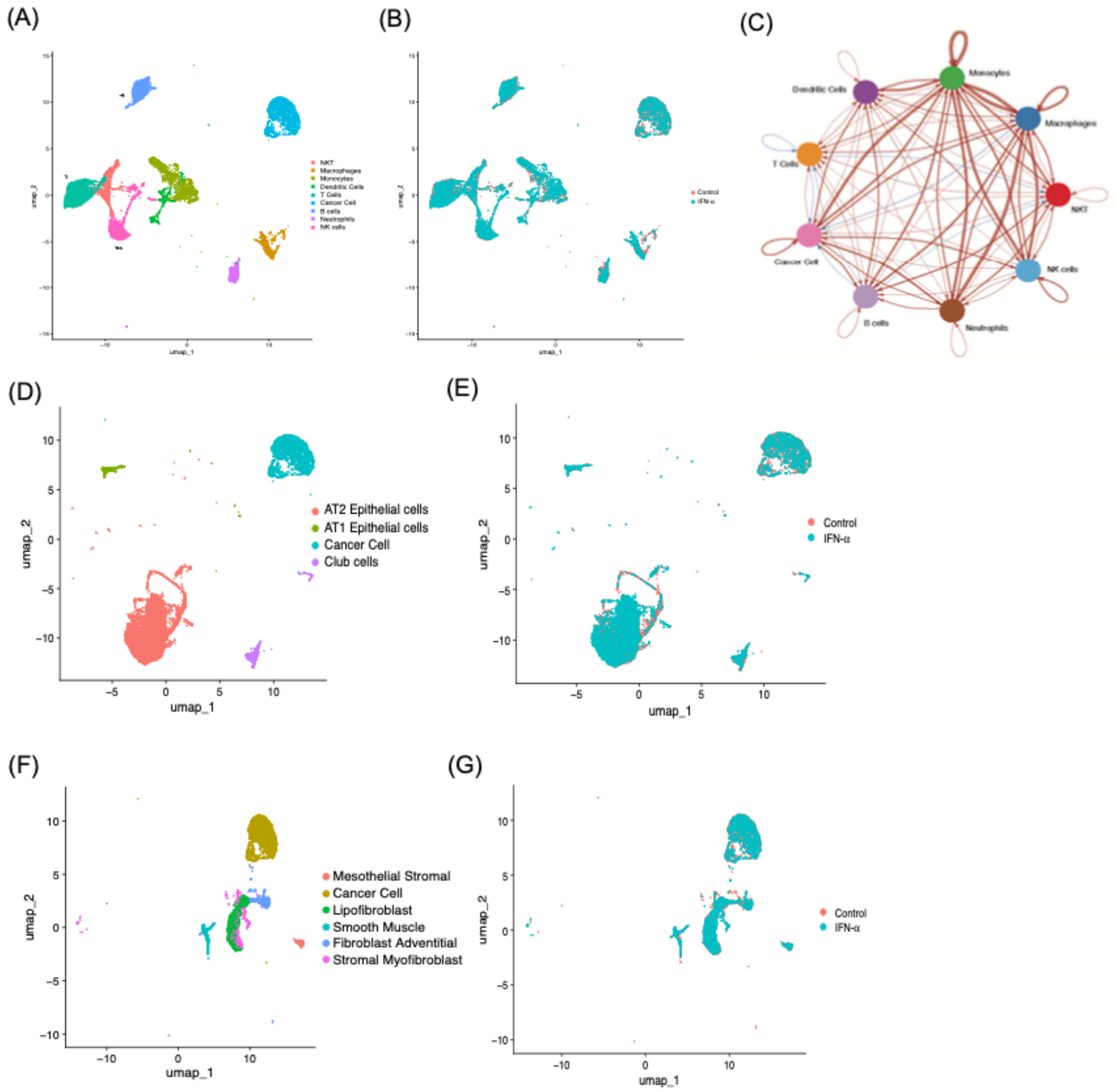


**Supp. Fig. 9. Clustering for CellChat Analysis.** UMAP plot of cell clusters (A) and treatment (B) examined for immune-cancer cell CellChat analysis. (C) Circle plot visualizing the directionality of Differential Interaction Score of Cancer Cells with immune cell components as based on Secreted ligands and Receptor interactions. UMAP plot of cell clusters (D) and treatment (E) examined for epithelial-cancer cell CellChat analysis. UMAP plot of cell clusters (F) and treatment (G) examined for epithelial-cancer cell CellChat analysis.

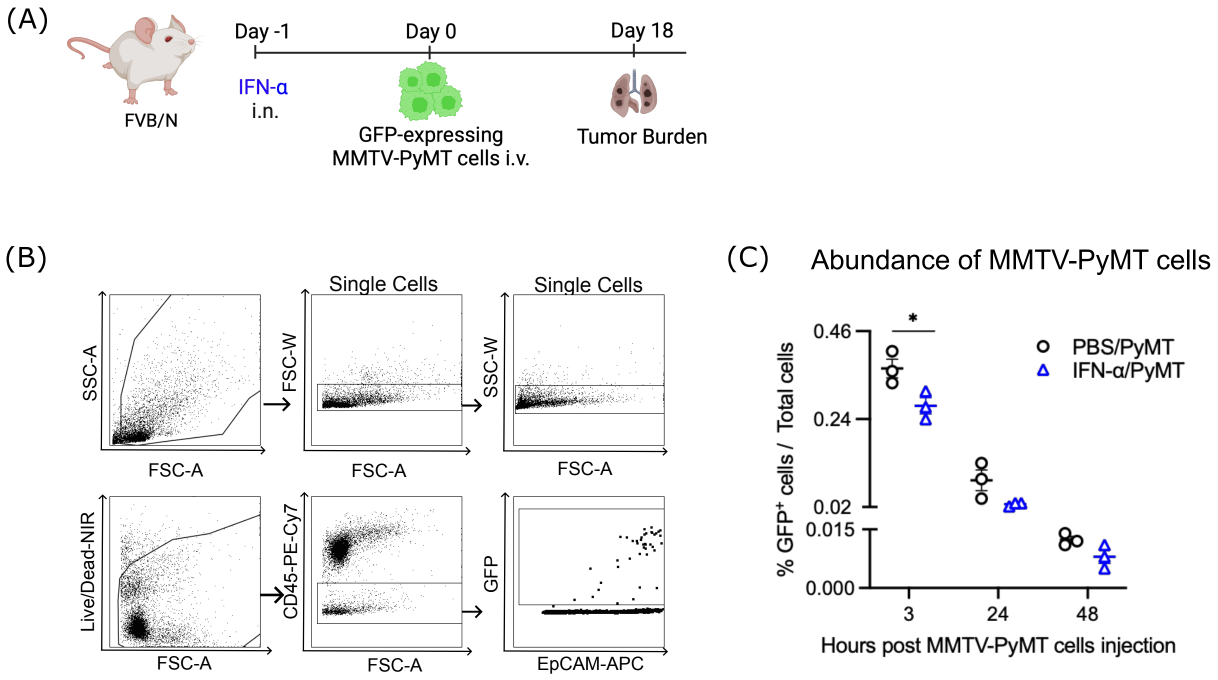


**Supp. Fig. 10. Intranasal exposure to IFN-α impairs homing of tumor cells to the lungs.** (A) FVB/N mice were intranasally exposed to 1ug of IFN-α. GFP-expressing MMTV-PyMT cells were injected i.v. 18h later. (B and C) Abundance of tumor cells in the lungs was quantified by flow cytometry, as shown in the gating strategy, at 3, 24 or 48 hours after tumor cell injection.

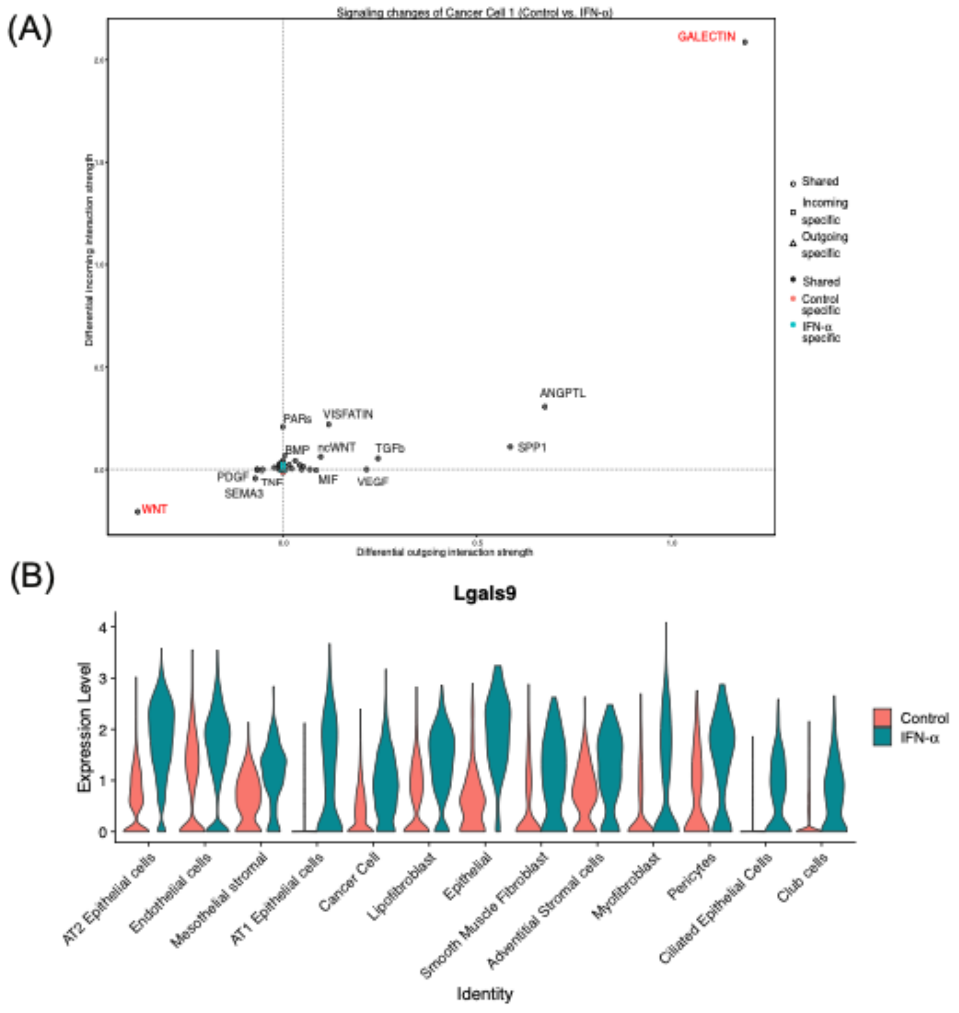


**Supp. Fig. 11. Changes in Cancer cell pathways.** (A) Plot of differential interaction strength scores for Secreted Ligand-Receptor interaction pathways in the Cancer Cell cluster, looking at predicted outgoing (secreted) and incoming (received) signals. (B) Violin plot of Galectin-9 (*Lgals9*) expression in non-immune cell clusters, split by treatment (Control-PBS, red or IFN-$\alpha$, blue).

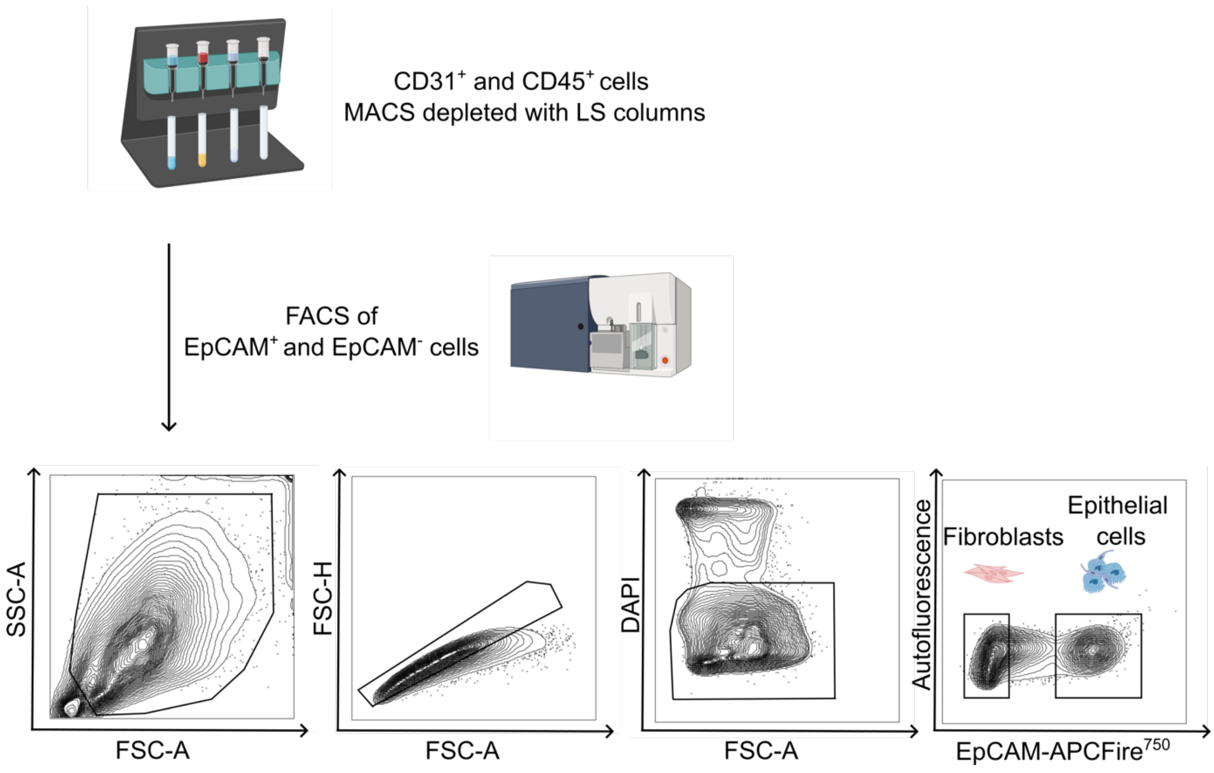


**Supp. Fig. 12. Gating strategy to isolate lung CD45$^-$CD31$^-$EpCAM$^+$ and CD45$^-$CD31$^-$EpCAM$^-$ cells after RSV or IFN-$\alpha$ administration.** C57BL/6J mice were infected i.n. with RSV, exposed to 1ug of IFN-$\alpha$ or mock infected (PBS). Lungs were harvested 18h later and were subjected to liberase digestion to obtain a single cell suspension. To enrich the populations of interest, lung CD45$^+$ and CD31$^+$ cells were depleted using LS columns. The remaining cells were stained for viability and EpCAM for further FACS sorting. Shown are representative flow cytometry plots from lung cells from a mock infected mouse.
